# Cardiac REDD1 alters glucose and fatty acid metabolic gene expression via an mTORC1-independent, PPARα-dependent mechanism and drives hypertrophic growth

**DOI:** 10.64898/2026.03.16.710895

**Authors:** Mason Wheeler, Jamie Renick, Roslyn Fawbush, Emily McAlpin, Shaunaci Stevens, Karthi Sreedevi, Junco Warren, Michael Dennis, Jessica Pfleger

## Abstract

**Background:** Regulated in development and DNA damage 1 (REDD1) is a highly inducible molecule that plays a role in numerous physiological and pathophysiological processes. It is a well-established negative regulator of mammalian target of rapamycin complex 1 (mTORC1), which is critical for maintaining elevated fatty acid-to-glucose oxidation ratio in the heart. In addition, REDD1 deletion results in hyperglycemia, suggesting that REDD1 is critical for tissue glucose metabolism. The role of REDD1 in regulating cardiac glucose and/or fatty acid metabolism in response to physiologic or pathophysiologic cues, however, remains unexplored.

**Methods:** Herein, we utilize AC16 cardiomyocytes with *REDD1* deletion, as well as mice with global or cardiomyocyte-specific deletion of *Redd1*, and their respective controls. We also subject these mice cardiac pressure overload using transverse aortic constriction (TAC) for 2 weeks or sham operation as a control. To examine the molecular regulators of glucose oxidation, we utilized qPCR and western blotting to evaluate pyruvate dehydrogenase (PDH) kinase (*PDK*) and phospho-PDH (pPDH) levels, respectively. We also directly measured PDH activity and glucose-driven cellular respiration. To investigate the complete REDD1-dependent transcriptome and metabolome, we performed RNA-sequencing (RNA-Seq) and untargeted metabolomics, respectively. To determine if the observed gene expression changes were dependent upon transcription factor peroxisome proliferator-activated receptor alpha (PPARα), we utilized an established pharmacologic PPARα inhibitor, GW6471. Here, we measured PPARα activity directly, as well as the expression of its target genes. In order to determine if our observed effects were mTORC1-dependent, we utilized mTORC1-specific inhibitor, everolimus. Finally, we measured cardiac hypertrophy using gravimetric analyses (heart weight (HW)-to-body weight (BW) or HW-to-tibia length (TL) ratios) and histological analyses of cardiomyocyte cross sectional area (CSA). We also measured mRNA and protein levels of pathological hypertrophic markers *Natriuretic Peptide B* (*Nppb)* and Cardiac Ankyrin Repeat Protein (CARP), respectively.

**Results:** Our data demonstrate that physiological levels of glucose induce REDD1 expression in cardiomyocytes. Further, we show that in cardiomyocytes or the hearts of mice with *REDD1* deletion, there is elevated *PDK4* expression, as well as increased levels of pPDH (S300 and/or S293) and reduced PDH activity. Interestingly, everolimus treatment has no effect on these alterations. *In vitro*, we also observe elevated glycolysis and glycolytic capacity, and reduced maximal respiratory capacity (MRC) in the presence of glucose. Interestingly, our RNA-Seq data reveals the upregulation of genes involved in fatty acid catabolism. Further, we demonstrate that PPARα activity is enhanced, and everolimus treatment also has no effect on this parameter. Additionally, we show that treatment of cardiomyocytes with GW6471 normalizes the expression of its target genes (*PDK4*, *ACSL1*) and levels of pPDH (S300), that are elevated in cells with *REDD1* deletion. Finally, we observe elevated REDD1 in the hearts of mice following TAC. Moreover, we show reduced HW/BW, HW/TL, cardiomyocyte CSA, and levels of cardiac *Nppb* and CARP in mice with cardiomyocyte *Redd1* deletion subjected to TAC versus controls also subjected to TAC. Importantly, TAC-induced reductions in cardiac *Pdk4* and pPDH (S293 and S300), are normalized to control levels in mice with *Redd1* deletion subjected to TAC.

**Conclusions:** Together, our findings suggest that physiological glucose-induced and pathological pressure overload-induced REDD1 is required for enhancing glucose oxidation and suppressing fatty acid oxidation in cardiomyocytes. In this way, REDD1 supports cardiac hypertrophic growth. We also outline a mechanism whereby REDD1 inhibits PPARα activity, thereby inhibiting the expression of its target genes, including *PDK4* and those involved in fatty acid oxidation. Finally, we demonstrate that these effects are independent of REDD1’s ability to inhibit mTORC1.

## Introduction

Metabolism is a tightly regulated network of interconnected pathways that regulate energy production, as well as other cellular processes. In the heart, fatty acids are the primary metabolic fuel^1^, while glucose and other metabolic substrates are utilized to a greater extent during various physiological (*i.e.*, insulin stimulation)^2^ and pathological (*i.e.*, pathological hypertrophic growth)^3^ conditions. Importantly, the complete molecular mechanisms underlying cardiac fuel preference under various conditions remain unclear.

Regulated in development and DNA damage 1 (REDD1) (also known as DNA damage inducible transcript 4 (DDIT4)) has been shown to be induced by both physiological stimuli, including glucose^4^, insulin^5, 6^, and exercise^7^, as well as pathological stress, including hypoxia^5^, insulin resistance^8, 9^, and aging^10^. REDD1 is also a well-established negative regulator of mammalian target of rapamycin complex 1 (mTORC1)^11^, which controls cellular growth and metabolism^12^. While the role of REDD1 has been well studied in other tissues, including the skeletal muscle^7, 13^ and retina^14, 15^, its role in the heart has not been thoroughly investigated.

Interestingly, work from Zhu *et al*. shows that in mice with cardiomyocyte-specific deletion of mTOR, the catalytic subunit of mTORC1, there is reduced cardiac palmitate oxidation and enhanced glucose oxidation, suggesting that mTOR is critical for maintaining the elevated fatty acid metabolism that is characteristic of the healthy heart^16^. Consistently, work from Dungan *et al*. demonstrates that in mice with global REDD1 deletion and hyperactive mTORC1 signaling, there is systemic glucose and insulin intolerance, suggesting defects in tissue glucose uptake and metabolism^13^. However, glucose oxidation was not examined in these mice. Thus, it was our hypothesis that REDD1 is a critical regulator of cardiomyocyte glucose and fatty metabolism.

Indeed, our findings demonstrate a role for cardiac REDD1 in controlling metabolic fuel preference, as well as the underlying molecular mechanism that controls this process. Namely, glucose-induced REDD1 enhances glucose and suppresses fatty acid metabolism via suppression of transcription factor, peroxisome proliferator activated receptor alpha (PPARα), and its target genes. Specifically, PPARα increases the expression of pyruvate dehydrogenase (PDH) kinase 4 (PDK4)^17, 18^ that phosphorylates and inhibits PDH, thereby inhibiting glucose oxidation^19–21^. In addition, PPARα increases the expression of fatty acid catabolic genes^17, 22, 23^. Moreover, we demonstrate that this process is mTORC1-independent. Finally, we examined this role for REDD1 in cardiac metabolic substrate preference in a model of cardiac pathology, namely pressure overload-induced cardiac hypertrophy. Notably, a shift from fatty acid to glucose oxidation supports early adaptive hypertrophic growth in this context^24–27^. Indeed, we find that REDD1 is required for the metabolic and structural remodeling that underlies cardiac hypertrophic growth.

These findings are noteworthy as they define REDD1 as a critical nodal regulator of cardiac glucose versus fatty acid utilization, uncover the mechanism by which this occurs, and reveal its essential role in cardiac pathology. In addition, it highlights a novel function for REDD1 that is independent of its role as an mTORC1 inhibitor. Identifying and understanding the functions of critical metabolic regulators, such as REDD1, during physiological and pathophysiological conditions is critical for the therapeutic targeting of these molecules and the development of novel disease therapies.

## Methods

### Cell culture

Human AC16 cardiomyocytes were obtained from Sigma-Aldrich (SCC109). AC16 cardiomyocytes with REDD1 deletion (AC16Δ*REDD1*) were generously provided by Drs. Shaunaci Stevens and Michael Dennis and generated as previously described^4, 28^. Cells were cultured in Dulbecco’s Modified Eagles Medium (DMEM)/Ham’s F-12 50/50 mix (containing 17.2 mM glucose, 1 mM pyruvate) (Corning), supplemented with 12.5% fetal bovine serum (FBS; Corning) and 1% penicillin/streptomycin (Gibco). Cells were used to passage 11. As indicated, cells were exposed to DMEM, no glucose (Gibco) supplemented with 1% penicillin/streptomycin, and treated with metabolic substrates and/or everolimus (Sigma-Aldrich-SML2282) or GW6471 (SelleckChem-S2798), as indicated.

Primary adult mouse cardiomyocytes were isolated from 10-14-week-old mice of the indicated genotypes using the simplified Langendorff-free method^29^. Briefly, mice were anesthetized, subjected to thoracotomy, and their hearts were flushed, clamped at the aorta, and excised. Following clearing, digestion was performed via collagenase buffer injection into the left ventricle (LV). Following digestion, the heart was gently teased apart and triturated. Stop solution containing 10% fetal bovine serum was added and the cells were passed through a 100 μm pore-size strainer. The cardiomyocytes were allowed to settle by gravity, while the non-myocytes were collected from the supernatant. Both myocyte and non-myocyte fractions were subjected to centrifugation and subjected to further downstream assays, as described.

### Animal Models

All animal procedures were performed in accordance with the National Institutes of Health *Guide for the Care and Use of Laboratory Animals*. All animal protocols were approved by the Institutional Animal Care and Use Committee of Virginia Tech (24-283). Mice with global REDD1 deletion (*Redd1^-/-^*) were generously provided by Dr. Michael Dennis, and were generated on the B6/129F1 background as previously described^30^. Mixed cohorts of 8-14-week-old male and female mice were used for experiments. Wild type (WT) mice (*Redd1^+/+^*) were used as controls.

Mice with cardiomyocyte-specific REDD1 deletion were generated by crossing *Redd1-*floxed (*Redd1*^Fl/Fl^) mice (generated on the C57BL/6 background and generously provided by Dr. David L. Williamson) to transgenic mice expressing Cre driven by the alpha myosin heavy chain promoter on the C57BL/6 background (*Redd1*^+/+, ⍺MHC-Cre^) (The Jackson Laboratory-Stock No: 011038). Mixed cohorts of 8-15-week-old male and female mice were used for experiments. *Redd1^+/+^*, *Redd1*^+/+, ⍺MHC-Cre^, and *Redd1*^Fl/Fl^ mice were used as controls.

### Transverse Aortic Constriction

Transverse aortic constriction (TAC) was performed as previously described^31^. Briefly, 10-12-week-old mice were anesthetized via inhalation of 3.0% isoflurane and intubated. The transverse thoracic aorta was dissected, and a surgical clip (calibrated to 0.08 mm) was placed between the innominate artery and the left common carotid artery and clamped around the aorta. Sham operations were performed using the same procedure without aortic constriction. The mice were then extubated, their chests closed, and subcutaneous injections of 0.1 mg/kg buprenorphine (immediately and day 3 post-operation) and 5 mg/kg ketoprofen (immediately and days 1 and 2 post-operation) were administered).

### Western blotting

20-30 µg of AC16 whole cell lysate, 7.5-15 µg of cardiac whole lysate, or 50 µg of AC16 or cardiac nuclear lysates were resolved using 4-20% polyacrylamide gel electrophoresis (PAGE) under denaturing conditions (Criterion Midi Protein Gel, Bio-Rad). Following transfer to nitrocellulose membranes, anti-REDD1 (ProteinTech-10638-1-AP), - pP70S6K (T389) (Cell Signaling-9205), -P70S6K (Cell Signaling-9202), -phospho-PDHA1 (S293) (Abcam-ab92696), -phospho-PDHA1 (S300) (Sigma-AP1064), -PDHA1 (Abcam-ab110330), - PPAR⍺ (Cayman Chemical-101710), -RXR⍺ (Cell Signaling-5388), -ACSL1 (Cell Signaling-4047), or -CARP (Santa Cruz-365056) antibodies were used at a dilution of 1:1000 in 3% bovine serum albumin (BSA; Fisher Scientific) in TBS-T. Signals were detected using anti-rabbit 680 or anti-mouse 800 secondary antibodies at a dilution of 1:10,000 in 3% BSA in TBS-T, washed, imaged using the Odyssey CLx Imaging System (LI-COR), and quantified using densitometry. Revert^TM^ 700 Total Protein Stain (LI-COR) was used to detect and quantify total protein on nitrocellulose membranes.

### Quantitative polymerase chain reaction (qPCR)

Total RNA was extracted using TRIzol reagent (ThermoFisher Scientific) as described by the manufacturer. 2 μg RNA was reverse transcribed to cDNA using the High-Capacity cDNA Reverse Transcription Kit (Applied Biosystems). qPCR was performed using TaqMan Gene Expression Assays (ThermoFisher Scientific) and the QuantStudio 3 Real-Time PCR System (ThermoFisher Scientific) for the following genes: 18S (Mm03928990_g1), *Redd1* (Mm00512504_g1), *PDK1* (Hs05380290_s1), *Pdk1* (Rn00587598_m1), *PDK2* (Hs04965351_m1), *Pdk2* (Rn00446679_m1), *PDK3* (Hs03878443_s1), *Pdk3* (Rn01424337_m1), *PDK4* (Hs01037712_m1), *Pdk4* (Rn00585577_m1), *PDP1* (Hs01081518_s1), *Pdp1* (Rn01437077_m1), *PDP2* (Hs01934174_s1), *Pdp2* (Mm02526496_s1), *ACSL1* (Hs00960561_m1), or *Nppb* (Mm01255770_g1).

### Mitochondrial stress test

AC16 or AC16Δ*REDD1* cardiomyocytes were plated at the indicated densities in 96-well XF Analyzer plates. 24 hours after plating, the medium was changed, as indicated, for 1 h. OCR and ECAR were measured using the XFe96 Analyzer (Agilent), as recommended by the manufacturer. Briefly, the medium of each well was injected with 1 μM oligomycin, the indicated dose of FCCP, and 1 μM antimycin A plus 1 μM rotenone at 18, 36, and 54 minutes after the start of measurements, respectively. Three readings were taken before and after the injection of each compound and the results plotted as pMol/min or mpH/min (y-axis) versus time (min) (x-axis).

### PDH activity assay

AC16 or AC16Δ*REDD1* cardiomyocytes were grown to ∼80% confluency and cultured in DMEM, no glucose supplemented with 5.5 mM glucose for 24 h. 15 million cells from each sample were subjected to mitochondrial isolation using the Mitochondria Isolation Kit (Sigma-Aldrich-MITOISO2), as described by the manufacturer. 20 µL of lysate was used in duplicate to measure PDH Activity (MAK183-Sigma-Aldrich).

### Echocardiography

10-14-week-old *Redd1*^Fl/Fl^ or *Redd1*^Fl/Fl, ⍺MHC-Cre^ mice were anesthetized using isoflurane and echocardiography was performed using the Vevo 2100 Imaging System (VisualSonics). For functional analyses, two-dimensional (2D) parasternal short-axis echocardiographic views were taken mid-ventricle at the level of the papillary muscles, below the mitral valve. In a plane bisecting the LV, motion (m)-mode measurements were used to calculate systolic and diastolic diameter (mm), systolic and diastolic volume (µL), stroke volume (µL), ejection fraction (%), fractional shortening (%), and cardiac output (mL/min). For aortic pressure measurements, 2D suprasternal echocardiographic views of the ascending aorta, aortic arch, and thoracic descending aorta were taken. Color and pulse wave (PW) Doppler was used to calculate mean and peak aortic pressure gradients (mmHg). Echocardiography was performed in a blinded fashion with animal genotypes disclosed upon completion of the studies.

### Gravimetric Analyses

Individual mouse body weight (BW), heart weight (HW) following excision, and tibia length (TL) following dissection, were measured. HW was divided by BW (HW/BW) or TL (HW/TL). Wet-to-dry lung weight ratio (Wet/Dry Lung Weight) was determined by measuring wet lung weight of each individual mouse immediately following excision. The lungs were then dried for 72 h at 60°C and re-weighed, corresponding to dry lung weight.

### Histological Staining

Heart tissue was fixed in 10% neutral buffered formalin, dehydrated, paraffin embedded, and sectioned (5 µm thickness). Following deparaffinization and rehydration, heart sections were stained with hematoxylin and eosin (H&E) or 594 dye-conjugated wheat germ agglutinin (WGA) (Invitrogen-W11263). Slides were mounted with Cytoseal (Epredia) or ProLong^TM^ Gold Antifade (Invitrogen).

### Blood Glucose Measurements

Whole blood was obtained via lateral saphenous venipuncture as previously described^32^. Briefly, the mice were restrained and the left hindleg immobilized to access the saphenous vein. A 27 g needle was used to draw blood. Whole blood was assessed using the OneTouch Verio Reflect blood glucometer.

### RNA-Sequencing

Standard RNA-Sequencing was performed by GENEWIZ from Azenta Life Sciences using the Illumina HiSeq 2x150bp platform. Sequence reads were trimmed and de-duplicated using fastp (v.0.23.1) and then aligned to the Mus musculus GRCm38 reference genome using the STAR Aligner (v.2.5.2b). Unique gene hit counts were determined using the Subread package-featureCounts (v.1.5.2). Differential expression analysis was then performed using DESeq2.

### Metabolomics

Untargeted metabolomic screening was performed using the Shimadzu GCMS-TQ8050 NX EI/CI/NCI mass spectrometry system, as described previously^33–36^. Briefly, freeze-clamped cardiac tissue samples were homogenized in an extraction solution consisting of 90% methanol/water and containing internal standards (d4-succinate and d4-myristic acid). All samples were dried and derivatized sequentially with O-methoxylamine hydrochloride in pyridine, followed by N-Trimethylsilyl-N-methyl trifluoroacetamide prior to GC-MS analysis. Bioinformatic analysis was conducted using MetaboAnalyst 6.0.

### PPAR⍺ Activity Assay

*In vitro,* AC16 or AC16Δ*REDD1* cardiomyocytes were grown to ∼80% confluency and cultured in DMEM, no glucose supplemented with 5.5 mM glucose for 24 hours and everolimus treatment, as indicated. *In vivo,* hearts from 10-15-week-old *Redd1*^Fl/Fl^ or *Redd1*^Fl/Fl, ⍺MHC-Cre^ mice were excised and the ventricles isolated. Cells or tissues were then subjected to nuclear extraction (Abcam-ab221978). 50 µg of lysate was used in duplicate to measure PPAR⍺ binding activity as described by the manufacturer (Abcam-ab133107).

### Statistics

Significant differences between the means of 2 sample groups were calculated with unpaired *t* tests (equal variance, 2 tailed), and 1- or 2-way ANOVA with Tukey post hoc testing was used for multiple comparisons. Values of *p*<0.05 were considered significant (\**p*<0.05, \*\**p*<0.01, \*\*\**p*<0.001, \*\*\*\**p*<0.0001).

## Results

### Glucose-induced REDD1 enhances cardiomyocyte PDH activity *in vitro*

We first confirmed that glucose increases cardiomyocyte REDD1 expression, as is described for other cell types^4, 14^. Indeed, REDD1 is significantly elevated in WT AC16 human cardiomyocytes cultured in hypoglycemic (2.5 mM glucose), normoglycemic (5.5 mM glucose), and hyperglycemic (25 mM glucose) conditions versus cell cultured without glucose (0 mM). Notably, these effects are not dose dependent, as REDD1 increased to a similar extent with all concentrations of glucose (**Supp Fig 1A-B**, **Fig 1A-B**). In addition, REDD1 deletion increases mTORC1 signaling, as demonstrated by an increase in the phosphorylation of P70S6 kinase (pP70S6K (T389)), confirming REDD1’s established function as an mTORC1 signaling inhibitor^11^. Importantly, this inhibition occurs at all tested concentrations of glucose (**Supp Fig 1A&C, Fig 1A&C**).

**Figure 1.**
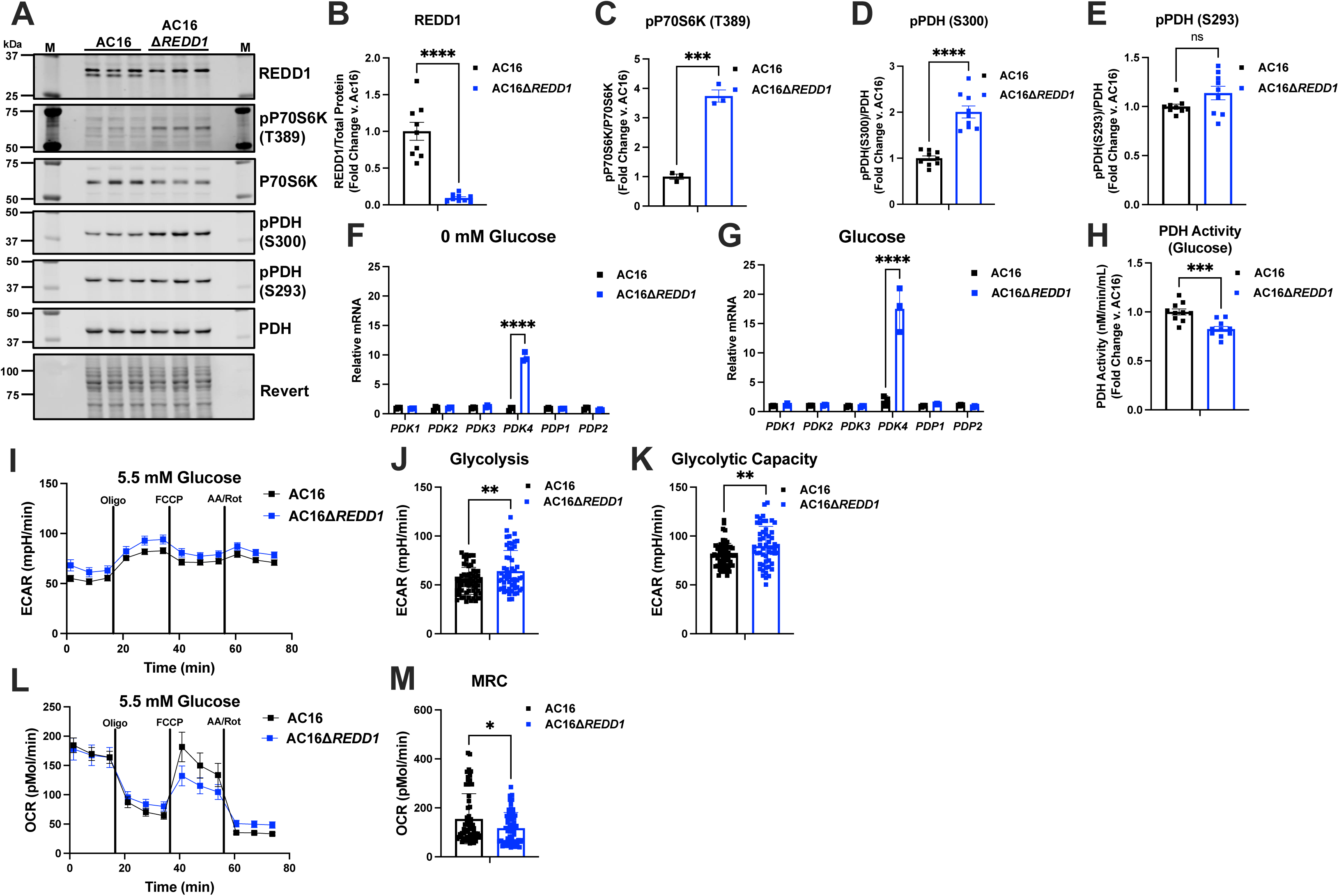
Glucose-induced REDD1 enhances cardiomyocyte PDH activity *in vitro.* **A-E.** AC16 and AC16Δ*REDD1* cardiomyocytes were cultured in DMEM, no glucose supplemented with 5.5 mM glucose for 24 hours and subjected to western blotting with the indicated antibodies. Signals were quantified with densitometry, normalized to total protein, total PDH, or total P70S6K, as indicated, and plotted. n=9,9 (REDD1, pPDH (S293), pPDH (S300)) and n=3,3 (pP70S6K (T389)), unpaired *t* test. AC16 and **F-G.** AC16Δ*REDD1* cardiomyocytes were cultured in **(F)** DMEM, no glucose, un-supplemented (0 mM Glucose) or **(G)** DMEM, no glucose supplemented with 5.5 mM glucose for 24 hours, total RNA extracted, and quantitative polymerase chain reaction performed for the indicated genes. n=3,3 (0 mM Glucose, 5.5 mM Glucose), 2-way ANOVA. **H.** AC16 and AC16Δ*REDD1* cardiomyocytes were cultured in DMEM, no glucose supplemented with 5.5 mM glucose for 24 hours, subjected to mitochondrial isolation, and PDH activity was measured. n=10,10, unpaired *t* test. **I-M.** AC16 and AC16Δ*REDD1* cardiomyocytes were plated at a density of 60,000 cells per well and subjected to mitochondrial stress tests as detailed in the methods section. The results are plotted as **(I)** extracellular acidification rate (ECAR) (mpH/min) versus time (minutes), with **(J)** glycolysis and **(K)** glycolytic capacity and **(L)** oxygen consumption (OCR) (pMol/min) versus time (minutes), with **(M)** maximal respiratory capacity. n=21,18 (ECAR), n=63,54 (Glycolysis, Glycolytic Capacity), n=21,19 (OCR), n=63,57 (MRC), unpaired *t* test. Error bars represent SEM. *p<0.05, **p<0.01, ***p<0.001, ****p<0.0001. M = marker.

In order to examine the role of REDD1 in regulating glucose oxidation, we evaluated the effects of REDD1 loss-of-function on the phosphorylation of pyruvate dehydrogenase (pPDH) at two known phosphorylation sites, serine 300 (S300) and serine 293 (S293). Phosphorylation at either of these sites is sufficient to inactivate the enzyme complex, preventing the decarboxylation of pyruvate and its conversion to acetyl CoA and entry into the tricarboxylic acid (TCA) cycle^21^. It is in this way that phosphorylation of PDH prevents the coupling of glycolysis to glucose oxidation, effectively preventing glucose oxidation^19–21^. We observe that REDD1 deletion increases pPDH (S300), demonstrating that REDD1 is required for the maintenance of unphosphorylated PDH during normoglycemic (5.5 mM glucose) (**Fig 1A&D**) and hypoglycemic (0 or 2.5 mM glucose) conditions, but not hyperglycemic (25 mM glucose) conditions (**Supp Fig 1A&D**). We also did not observe any alterations in pPDH (S293) (**Fig 1A&E, Supp Fig 1A**).

We also examined levels of the PDH kinases (PDKs) that phosphorylate and PDH phosphatases (PDPs) that dephosphorylate PDH^20, 21^. These data demonstrate that in AC16s with REDD1 deletion, there is a selective increase in *PDK4* versus WT AC16s with no alterations in *PDK1-3* or *PDP1* or *2*. This occurs in cells cultured in 0 or 5.5 mM glucose (**Fig 1F&G**). Consistent with elevated PDK4 and pPDH (S300), we observe reduced PDH activity in AC16s with REDD1 deletion versus WT AC16s (**Fig 1H**). Thus, REDD1 is required for the suppression of PDK4 to maintain PDH activity, and likely glucose oxidation, during hypoglycemic and normoglycemic conditions.

If pyruvate is not converted into acetyl CoA, one of its fates is conversion to lactate and extrusion from the cell^37, 38^. Consistently, we observe elevated extracellular acidification rates (ECAR), glycolysis, and glycolytic capacity in AC16s with REDD1 deletion versus WT controls cultured in normoglycemic conditions (5.5 mM glucose) (**Fig 1I-K**). Finally, we also observe a reduction in maximal respiratory capacity (MRC) in AC16s with REDD1 deletion versus WT controls cultured in normoglycemic conditions (5.5 mM glucose). Importantly, we do not observe any significant alterations in basal oxygen consumption rate (OCR), ATP-linked OCR, proton leak, or non-mitochondrial respiration (**Fig 1L-M**). These findings are consistent with our previously published data demonstrating a positive correlation of MRC with reducing or enhancing glucose oxidation via the overexpression of PDKs or inhibition of PDKs with dichloroacetate (DCA), respectively^39^. This is intuitive as the manipulation of glucose oxidation will not alter baseline energetics in the heart, but deficiencies are revealed under conditions of increased metabolic demand^39^. Of note, we do not observe a reserve respiratory capacity (RRC) in AC16s cultured in amino acids (0 mM glucose), 5.5 mM glucose, or the combination glucose (5.5 mM)+palmitate, but we do observe a RRC when cells are cultured in the presence of palmitate, indicating a more adult metabolic phenotype of these cells (**Supp Fig 2A**)^1, 39^. Cell number and FCCP concentrations for these experiments were optimized at 60,000 cells per well and 1 μM, respectively (**Supp Fig 2B-C**).

**Figure 2.**
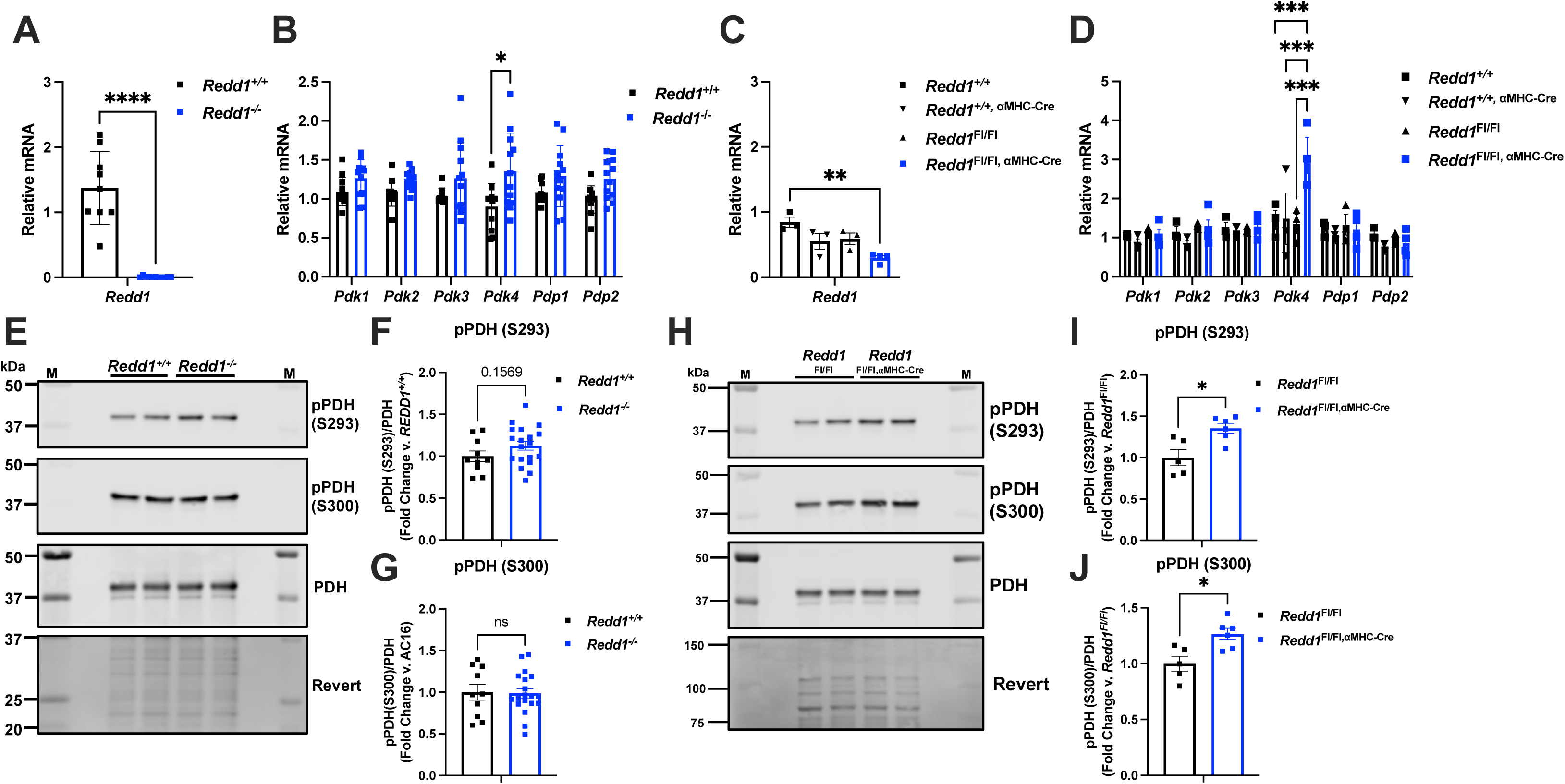
REDD1 alters cardiomyocyte PDH regulatory molecules *in vivo.* **A-D.** Hearts of adult (8-14-week-old) male and female mice of the indicated genotypes were harvested and subjected to total RNA extraction and qPCR for the indicated genes. **(A-B)** n=9,11 (*Redd1)*, n=12,12 (*Pdk1, Pdk2, Pdp1*), n=10,12 (*Pdk3*), and n=11,12 (*Pdk4, Pdp2*), unpaired *t* test, 2-way ANOVA. **(C-D)** n=3,3,3,4 (*Redd1, Pdk1, Pdk2, Pdk3, Pdp1, Pdp2*) and n=3,3,3,3 (*Pdk4*), 1-way ANOVA, 2-way ANOVA. **E-J.** Hearts of adult (8-14-week-old) male and female mice of the indicated genotypes were harvested, lysed, and subjected to western blotting with the indicated antibodies. Signals were quantified with densitometry, normalized to total PDH, and plotted. **(E-G)** n=10,19 (pPDH (Ser293), pPDH (Ser300)), unpaired *t* test. **(H-J)** n=5,6 (pPDH (Ser293), pPDH (Ser300)), unpaired *t* test. Error bars represent SEM. *p<0.05, **p<0.01, ***p<0.001, ****p<0.0001. M = marker.

Together these data demonstrate that under physiological, normoglycemic conditions, elevated REDD1 is required for the suppression of PDK4 and pPDH (S300) levels, contributing to elevated PDH activity and MRC, which are required for glucose oxidation^19–21^.

### REDD1 alters cardiomyocyte PDH regulatory molecules *in vivo*

To test the effects of cardiomyocyte REDD1 loss-of-function *in vivo*, we utilized mice with global deletion of REDD1 (*Redd1*^-/-^), as well as generated novel mice with cardiomyocyte-specific deletion of REDD1 (*Redd1*^Fl/Fl,αMHC-Cre^). Our data demonstrate a complete loss of REDD1 expression and elevated *Pdk4* expression levels in our *Redd1*^-/-^ mice versus WT controls, with no significant alterations in *Pdk1-3* or *Pdp1* or *2* expression (**Fig 2A-B**). In the whole hearts from our mice with REDD1 cardiomyocyte-specific deletion, we observe a 55% reduction in *Redd1* expression levels versus combined controls (**Fig 2C**). Since REDD1 is ubiquitously expressed^40^, we isolated cardiomyocytes and non-myocytes from the hearts of these mice and demonstrate a loss of *Redd1* expression in the *Redd1*^Fl/Fl,αMHC-Cre^ mice versus controls, while we observe no significant differences in the non-myocyte fraction (**Supp Fig 3A-B**). Importantly, we also observe a selective increase in *Pdk4* expression in the hearts of *Redd1*^Fl/Fl,αMHC-Cre^ mice versus controls, with no significant alterations in *Pdk1-3* or *Pdp1* or *2* expression (**Fig 2D**). Importantly, these increases in *Pdk4* expression translate into a trending increase in pPDH (S293) (**Fig 2E-G**), as well as significant increases in pPDH (S293 and S300) in the mice with REDD1 global or cardiomyocyte-specific deletion, respectively (**Fig 2 H-J**). Notably, there are no significant differences in cardiac systolic function (**Supp Fig 4A-C; Supp Table 1**), cardiac size (**Supp Fig 4D-G; Supp Table 1**) or morphology (**Supp Fig 4H**), or fasted blood glucose (**Supp Fig 4I**) in the *Redd1*^Fl/Fl,αMHC-Cre^ mice versus *Redd1*^Fl/Fl^ controls. Consistently, Stevens *et al.* show no alterations in systolic function in the hearts of mice with global REDD1 deletion^41^. Together, these data demonstrate that under physiological conditions *in vivo*, REDD1 is required for the suppression of PDK4 and pPDH levels, which are required for glucose oxidation^19–21^.

**Figure 3.**
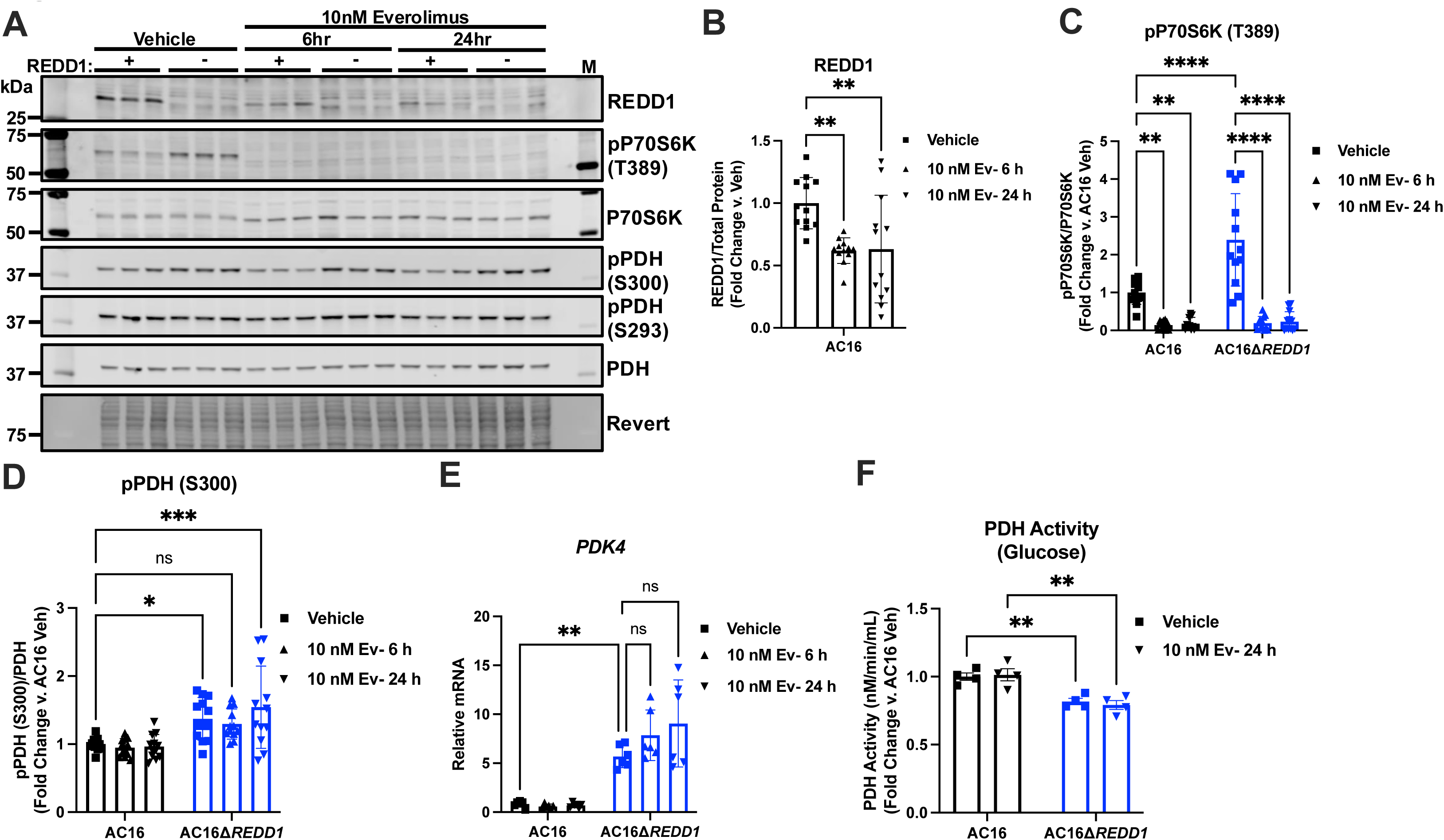
REDD1’s control of PDH activity is mTORC1-independent. AC16 and AC16Δ*REDD1* cardiomyocytes were cultured in DMEM, no glucose supplemented with 5.5 mM glucose for 24 hours and vehicle or 10 nM Everolimus treatment for 6 or 24 hours. **A-D.** The cardiomyocytes were subjected to western blotting with the indicated antibodies. Signals were quantified with densitometry, normalized to total protein, total PDH, or total P70S6K, as indicated, and plotted. n=12,12,12 (REDD1), n=12,12,12,12,12,12 (pP70S6K (T389) and pPDH (S300)), 1-way ANOVA, 2-way ANOVA. **E.** The cardiomyocytes were subjected to total RNA extraction and qPCR for *PDK4*. n=6,6,6,6,6,6, 2-way ANOVA. **F.** The cardiomyocytes were subjected to mitochondrial isolation, and PDH activity was measured. n=4,4, 2-way ANOVA. Error bars represent SEM. *p<0.05, **p<0.01, ***p<0.001, ****p<0.0001. M = marker.

**Figure 4.**
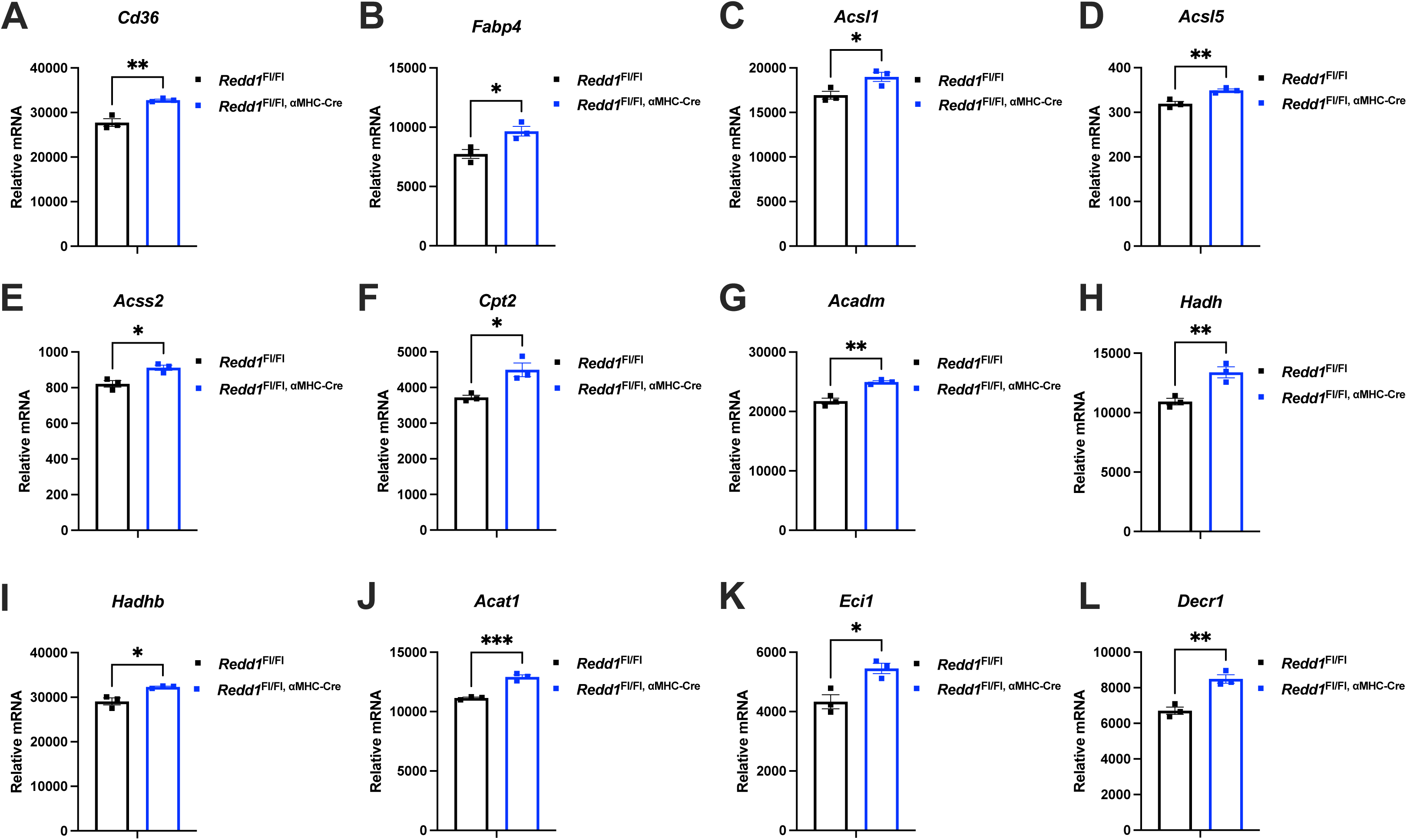
REDD1 inhibits the expression of genes involved in fatty acid catabolism. **A-L.** Hearts of adult (12-week-old) male *Redd1*^Fl/Fl^ and *Redd1*^Fl/Fl, ⍺MHC-Cre^ mice were excised and subjected to RNA-sequencing. Genes involved in fatty acid catabolism are graphed. n=3,3 (*Cd36*, *Fabp4*, *Acsl1*, *Acsl5*, *Acss2*, *Cpt2*, *Acadm*, *Hadh*, *Hadhb*, *Decr1*, *Eci1*, *Acat1*), unpaired *t* test. Error bars represent SEM. *p<0.05, **p<0.01, ***p<0.001.

**Table 1.** Gene Ontological Analysis. Gene ontological analysis was performed using The Database for Annotation, Visualization, and Integrated Discovery (DAVID) for the differentially expressed genes between *Redd1*^Fl/Fl^ and *Redd1*^Fl/Fl, ⍺MHC-Cre^ mice as assessed by RNA sequencing. Top results and their respective counts and p-values are listed.

| Term | Count | p Value |
| --- | --- | --- |
| Fatty Acid Metabolic Process | 16 | 1.8E-4 |
| Fatty Acid Beta-Oxidation | 8 | 5.6E-4 |
| Lipid Biosynthetic Process | 5 | 3.7E-3 |
| Mitochondrial Electron Transport | 5 | 4.2E-3 |
| BCAA Catabolic Process | 4 | 4.5E-3 |
| Transcription, DNA-Templated | 9 | 5.5E-3 |
| Lipid Metabolic Process | 31 | 6.2E-3 |
| Angiogenesis | 16 | 6.6E-3 |
| Endocytosis | 13 | 8.6E-3 |
| Regulation of Cell Migration | 9 | 8.9E-3 |
| ATP Synthesis Couple to Electron Transport | 3 | 1.2E-2 |
| Regulation of Mito Membrane Potential | 5 | 1.3E-2 |
| Lipid Transport | 11 | 1.3E-2 |

### REDD1’s control of PDH activity is mTORC1-independent

Since REDD1 is an established negative regulator of mTORC1 signaling^11^, we investigated whether or not the effects of REDD1 on glucose oxidation were mTORC1 dependent. To do so, we utilized a selective mTORC1 inhibitor, everolimus^42^. Our data show that REDD1 levels are reduced with both 6 or 24 hour everolimus treatment (**Fig 3A-B**), which is consistent with reports demonstrating mTORC1’s control of REDD1 expression as part of a negative feedback loop^43^. In addition, both basal pP70S6K (T389) levels, as well as those that are enhanced as a result of REDD1 deletion, are abolished with 6 or 24 hour everolimus treatment, demonstrating the efficacy of the inhibitor (**Fig 3A&C**). Interestingly, we find that enhanced pPDH (S300) levels, as observed in cardiomyocytes with REDD1 deletion, are not normalized to WT levels with 6 or 24 hour everolimus treatment (**Fig 3A&D**). Consistently, elevated *PDK4* mRNA levels, as observed in cardiomyocytes with REDD1 deletion, are also not normalized to WT levels with 6 or 24 hour everolimus treatment (**Fig 3E**). Finally, PDH activity, which is reduced in cardiomyocytes with REDD1 deletion, is not normalized to WT levels with 24 hour everolimus treatment (**Fig 3F**). Together, these data demonstrate that REDD1-mediated inhibition of PDK4 expression and PDH phosphorylation, and REDD1-dependent PDH activity, which are required for glucose oxidation^19–21^, are mTORC1-independent.

### REDD1 inhibits the expression of genes involved in fatty acid catabolism

Since REDD1 controls the *PDK4* mRNA levels, we examined the complete set of transcripts regulated by REDD1. To do so, we performed RNA sequencing (RNA-Seq) in mice with cardiomyocyte-specific deletion of REDD1 (*Redd1*^Fl/Fl,αMHC-Cre^) or controls (*Redd1*^Fl/Fl^). Our gene ontology (GO) analysis shows ‘Fatty Acid Metabolic Process’ and ‘Fatty Acid Beta-Oxidation’ as top terms among others (**Table 1**). Similarly, KEGG pathway analysis uncovers ‘Fatty Acid Metabolism’, ‘Fatty Acid Degradation’, ‘Oxidative Metabolism’, and ‘Pyruvate Metabolism’ as top altered pathways among others (**Table 2**). Interestingly, when we specifically interrogated the mRNA levels of genes involved in fatty acid catabolism, we see an upregulation of those involved in cellular fatty acid uptake (*Cd36*) (**Fig 4A**), transport (*Fabp4*) (**Fig 4B**), conversion to fatty acyl-CoAs (*Acsl1*, *Acsl5*, *Acss2*) (**Fig 4C-E**), mitochondrial uptake (*Cpt2*) (**Fig 4F**), as well as beta-oxidation of saturated (*Acadm*, *Hadh*, *Hadhb*, *Acat1*) (**Fig 4G-J**) and unsaturated (*Eci1*, *Decr1*) (**Fig 4K-L**) fatty acids^17,22^. These data demonstrate that REDD1 negatively regulates the expression of genes involved in fatty acid catabolism. This, along with our data showing that REDD1 negatively regulates PDK4 expression and pPDH, and positively regulates PDH activity, suggests that cardiomyocyte REDD1 suppresses fatty acid, while activating glucose metabolism.

**Table 2.** KEGG Pathway Analysis. KEGG pathway enrichment analysis was performed using The Database for Annotation, Visualization, and Integrated Discovery (DAVID) for the differentially expressed genes between *Redd1*^Fl/Fl^ and *Redd1*^Fl/Fl, ⍺MHC-Cre^ mice as assessed by RNA sequencing. Top results and their respective counts and p-values are listed.

| Term | Count | p Value |
| --- | --- | --- |
| Valine, Leucine, Isoleucine Degradation | 11 | 1.6E-6 |
| Fatty Acid Metabolism | 9 | 1.8E-4 |
| Fatty Acid Degradation | 8 | 3.5E-4 |
| Diabetic Cardiomyopathy | 16 | 3.8E-4 |
| Metabolic Pathways | 60 | 3.0E-3 |
| Thermogenesis | 14 | 7.0E-3 |
| Propanoate Metabolism | 5 | 7.9E-3 |
| Oxidative Phosphorylation | 10 | 8.5E-3 |
| Efferocytosis | 11 | 8.7E-3 |
| Adherens Junctions | 8 | 9.5E-3 |
| Lipoic Acid Metabolism | 4 | 1.2E-2 |
| Chemical Carcinogenesis- ROS | 13 | 1.3E-2 |
| Alzheimer Disease | 18 | 2.1E-2 |
| Lysine Degradation | 6 | 2.4E-2 |
| Pyruvate Metabolism | 5 | 2.6E-2 |
| PPAR Signaling Pathway | 7 | 2.7E-2 |

### REDD1 inhibits PPARα activity *in vitro* and *in vivo* and is mTORC1-independent

We hypothesized that REDD1 is suppressing fatty acid and activating glucose metabolism at the transcriptional level. Specifically, we investigated transcription factor peroxisome proliferator-activated receptor alpha (PPARα), as it is a critical positive regulator of cardiac fatty acid metabolism and targets many of the genes uncovered in our RNA-Seq data (**Fig 4**)^17, 22, 23^. Indeed, ‘PPAR signaling pathway’ is a highly represented pathway in our KEGG pathway analysis (**Table 2**). In addition, PPARα has been shown to transcriptionally target *PDK4*^17, 18, 22^. Indeed, we observe a significant increase in PPARα activity in nuclei isolated from cardiomyocytes with REDD1 deletion versus controls **(Fig 5A)**. Importantly, this increase in activity is not normalized with everolimus treatment, suggesting that REDD1’s inhibition of PPARα activity is mTORC1-independent (**Fig 5A**). It is important to note that in WT AC16 cells, everolimus treatment does increase PPARα activity (**Fig 5A**), as mTORC1 has been well established to inhibit its activity via promoting nuclear accumulation of nuclear receptor corepressor 1 (nCoR1)^44^. Notably, there are no alterations in the expression levels of PPARα or its binding partner retinoid X receptor alpha (RXRα) in cardiomyocytes with REDD1 deletion and/or with everolimus treatment (**Fig 5B-D**).

**Figure 5.**
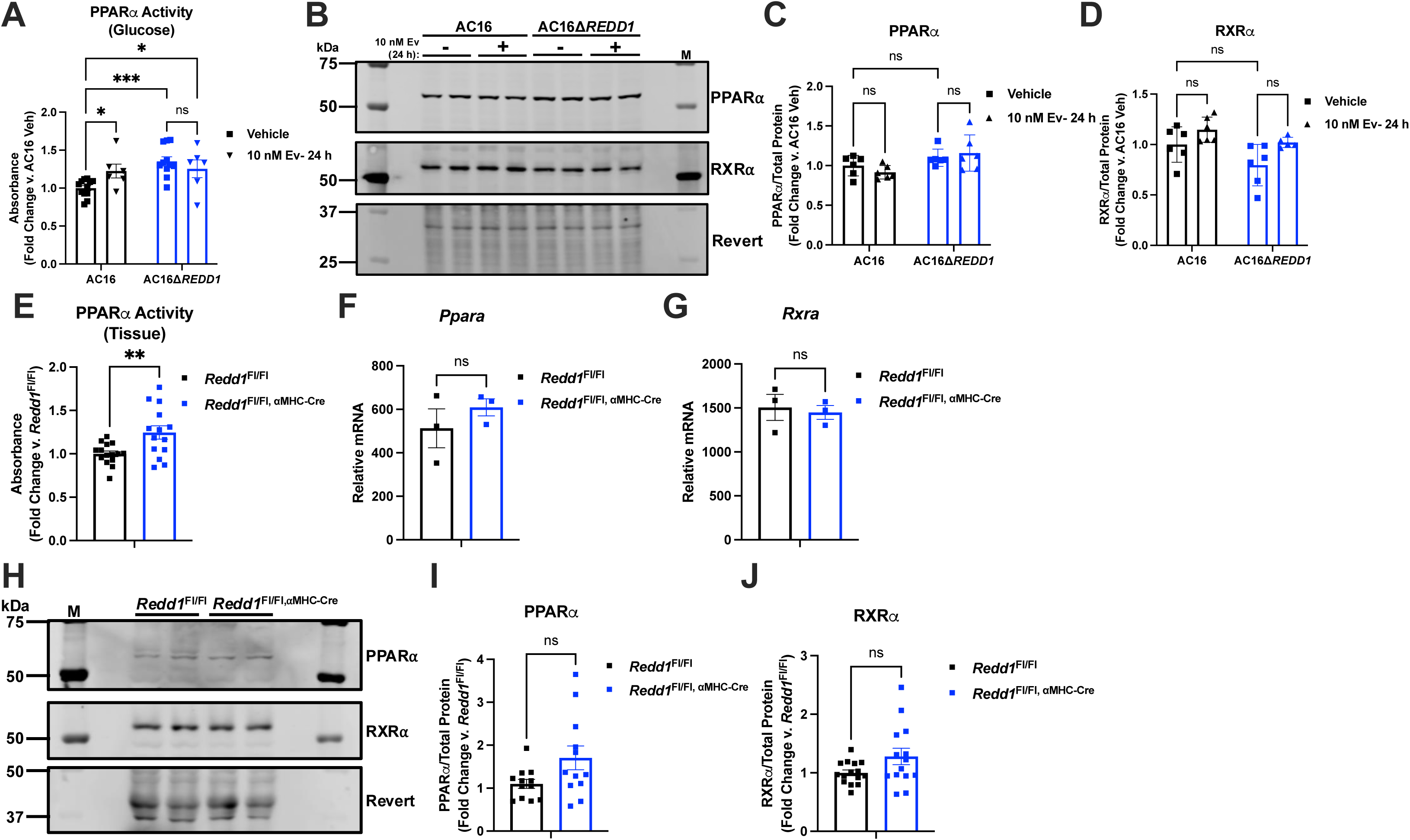
REDD1 inhibits PPAR⍺ activity *in vitro* and *in vivo* and is mTORC1-independent. **A-D.** AC16 and AC16Δ*REDD1* cardiomyocytes were cultured in DMEM, no glucose supplemented with 5.5 mM glucose and vehicle or 10 nM Everolimus treatment for 24 hours, subjected to nuclear extraction, assayed for **(A)** PPAR⍺ activity or subjected to **(B-D)** western blotting for the indicated antibodies. Signals were quantified with densitometry, normalized to total protein, and plotted. n=12,6,11,6 (PPAR⍺ activity), n=6,6,6,6 (PPAR⍺), n=6,6,6,5 (RXR⍺), 2-way ANOVA. **E, H-J.** Hearts of adult (10-15-week-old) male and female *Redd1*^Fl/Fl^ and *Redd1*^Fl/Fl, ⍺MHC-Cre^ mice were excised, subjected to nuclear extraction, assayed for **(E)** PPAR⍺ activity or subjected to **(H-J)** western blotting for the indicated antibodies. Signals were quantified with densitometry, normalized to total protein, and plotted. n=15,14 (PPAR⍺ activity, RXR⍺), n=12,12 (PPAR⍺), unpaired *t* test. **F-G.** Hearts of adult (12-week-old) male *Redd1*^Fl/Fl^ and *Redd1*^Fl/Fl, ⍺MHC-Cre^ mice were excised and subjected to RNA-sequencing. n=3,3 (*Ppara*, *Rxra*), unpaired *t* test. Error bars represent SEM. *p<0.05, **p<0.01, ***p<0.001. M = marker.

These findings are also confirmed *in vivo*, as mice with cardiomyocyte-specific deletion of REDD1 (*Redd1*^Fl/Fl,αMHC-Cre^) had elevated PPARα activity versus controls (*Redd1*^Fl/Fl^) (**Fig 5E**). Moreover, there were no significant alterations in the expression of PPARα or RXRα at the mRNA (**Fig 5F-G**) or protein (**Fig 5H-J**) levels. Since both PPARα and PDK4 activities are regulated by metabolites, namely PPARα is activated by fatty acids^45^ and PDK4 is inhibited by pyruvate and activated by acetyl CoA^21^, we examined the cardiac metabolomes of mice with global REDD1 deletion (*Redd1*^-/-^) or cardiomyocyte-specific REDD1 deletion (*Redd1*^Fl/Fl,αMHC-Cre^) and their controls (*Redd1*^+/+^ or *Redd1*^Fl/Fl^, respectively). Our findings demonstrate no significant differences in the tested metabolites from the hearts of either *Redd1*^-/-^ (**Supp Fig 5A**) or *Redd1*^Fl/Fl,αMHC-Cre^ (**Supp Fig 5B**) mice versus controls, suggesting that the effects of REDD1 on PPARα or PDH activities are not due to changes in metabolism. Taken together, these data demonstrate that REDD1 inhibits PPARα activity in an mTORC1-independent manner.

### REDD1’s control of glucose and fatty acid metabolic genes is PPARα-dependent

Next, we examined whether REDD1-mediated suppression of fatty acid catabolic genes and *PDK4* requires PPARα. To do so, we utilized a well-established selective inhibitor of PPARα activity, GW6471, which has been shown to disrupt its binding to coactivators and enhance its binding to corepressors^46, 47^. Indeed, our data show that the elevated PDK4 mRNA levels that we observe in the REDD1-deleted AC16s are partially normalized with GW6471 (**Fig 6A**). Consistently, the increase in pPDH (S300) that we observe in the REDD1-deleted AC16s was also partially normalized with GW6471 (**Fig 6C-D**). In addition, we confirm that *ACSL1* mRNA is increased in the AC16s with REDD1 deletion (**Fig 6B**), as is seen in the hearts of our mice with cardiomyocyte-specific REDD1 deletion (**Fig 4C**). This increase in ACSL1 is also seen at the protein level (**Fig 6C&E**) Moreover, these increases in ACSL1 were partially normalized with GW6471 (**Fig 6B, C&E**). Together, these data suggest that REDD1’s control of glucose oxidation and inhibition of fatty acid oxidation is at least partially dependent on PPARα activity.

**Figure 6.**
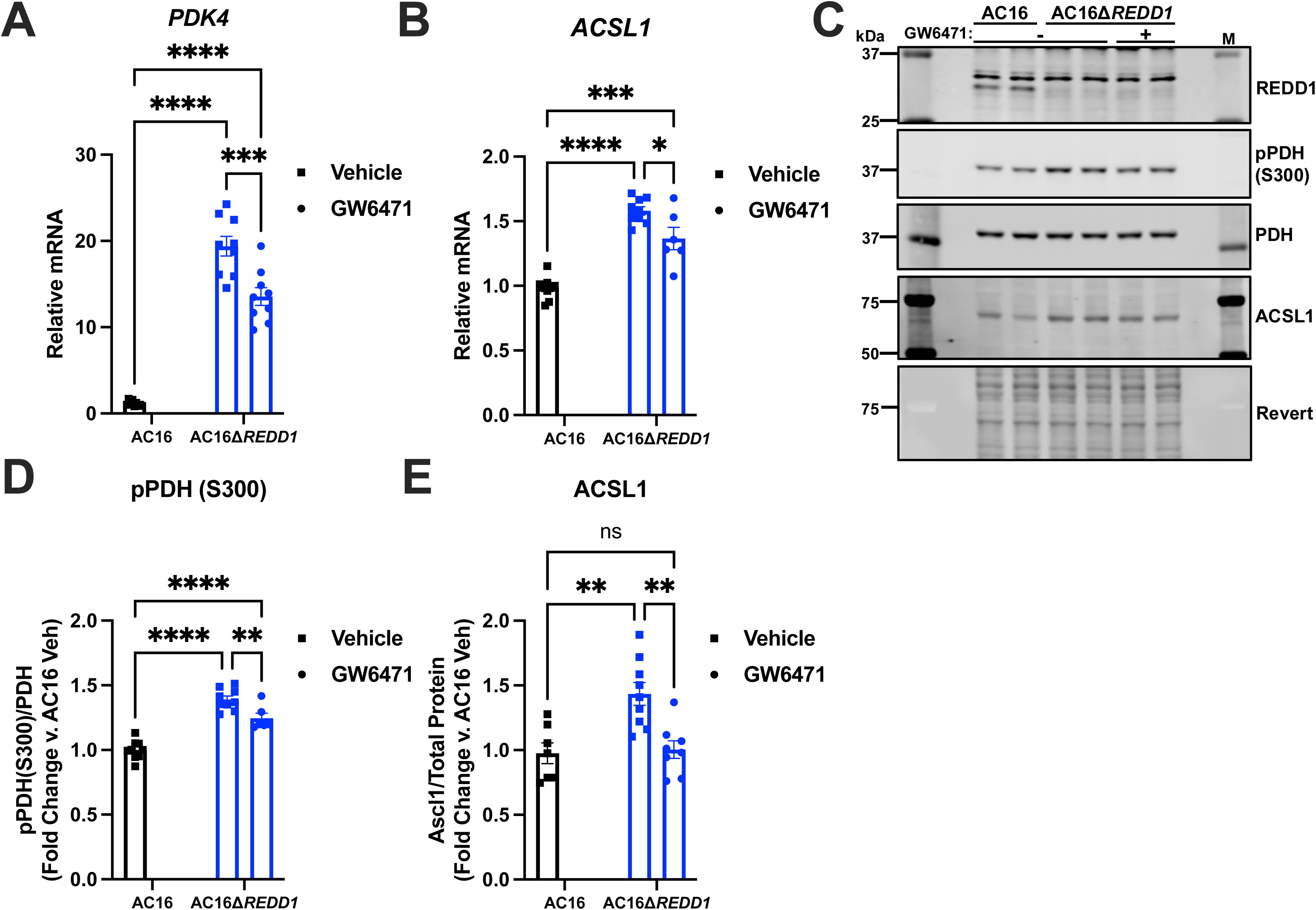
REDD1’s control of glucose and fatty acid metabolic genes is PPAR⍺-dependent. AC16 and AC16Δ*REDD1* cardiomyocytes were cultured in DMEM, no glucose supplemented with 5.5 mM glucose and vehicle or 0.1 µM GW6471 treatment for 24 hours. **A-B.** Total RNA was extracted and qPCR was performed for the indicated genes. n=9,9,9 (*PDK4*), n=9,9,6 (*ACSL1*), 2-way ANOVA. **C-E.** Cardiomyocytes were harvested and subjected to western blotting with the indicated antibodies. Signals were quantified with densitometry, normalized to total protein or PDH as indicated, and plotted. n=9,9,6 (pPDH (S300)), n=7,9,8 (ACSL1), 2-way ANOVA. Error bars represent SEM. *p<0.05, **p<0.01, ***p<0.001, ****p<0.0001. M = marker.

### Cardiac REDD1 is required for metabolic and structural remodeling in pathological hypertrophy

Since REDD1 alters gene expression consistent with elevated glucose and reduced fatty acid oxidation, we examined its role in pathological cardiac hypertrophy, a process that requires this shift in metabolic substrate utilization (namely, enhanced glucose and suppressed fatty acid oxidation)^24–27^. To do so, we subjected mice with cardiomyocyte-specific deletion of REDD1 (*Redd1*^Fl/Fl,αMHC-Cre^) or controls (*Redd1*^Fl/Fl^) to transverse aortic constriction (TAC) surgery for two weeks. Sham operation was used as a surgical control. Importantly, mean and peak aortic pressures were significantly elevated in the TAC versus sham-operated mice, but did not differ between the *Redd1*^Fl/Fl^ and *Redd1*^Fl/Fl,αMHC-Cre^ mice, indicating success of the operation and its consistency between experimental groups, respectively (**Supp Fig 6A-B**). Using this model, we observe increased *Redd1* mRNA levels in the hearts of *Redd1*^Fl/Fl^ control mice subjected to TAC versus sham operation (**Fig 7A**). Moreover, we observe pathological cardiac hypertrophy in *Redd1*^Fl/Fl^ control mice subjected to TAC versus sham operation, as assessed by increased heart weight-to-body weight (**Fig 7B**) and heart weight-to-tibia length ratios (**Fig 7C**), increased myocyte cross-sectional area (**Fig 7D-E**), and increased *Natriuretic Peptide B* (*Nppb)* mRNA **(Fig 7F)** and Cardiac Ankyrin Repeat Protein (CARP) (**Fig 7G-H)** levels. *Nppb* and CARP are well established markers of pathological hypertrophy^48^. Notably, this pathological cardiac hypertrophy is significantly reduced in mice with cardiomyocyte-specific REDD1 deletion (*Redd1*^Fl/Fl,αMHC-Cre^ + TAC versus *Redd1*^Fl/Fl^ + TAC) (**Fig 7 B-H**). It is also important to note that there was no change in wet-to-dry lung weight ratio in TAC versus sham operated mice of either genotype (*Redd1*^Fl/Fl^ or *Redd1*^Fl/Fl,αMHC-Cre^), indicating a lack of congestive heart failure in all groups ^49^(**Supp Fig 6C**).

**Figure 7.**
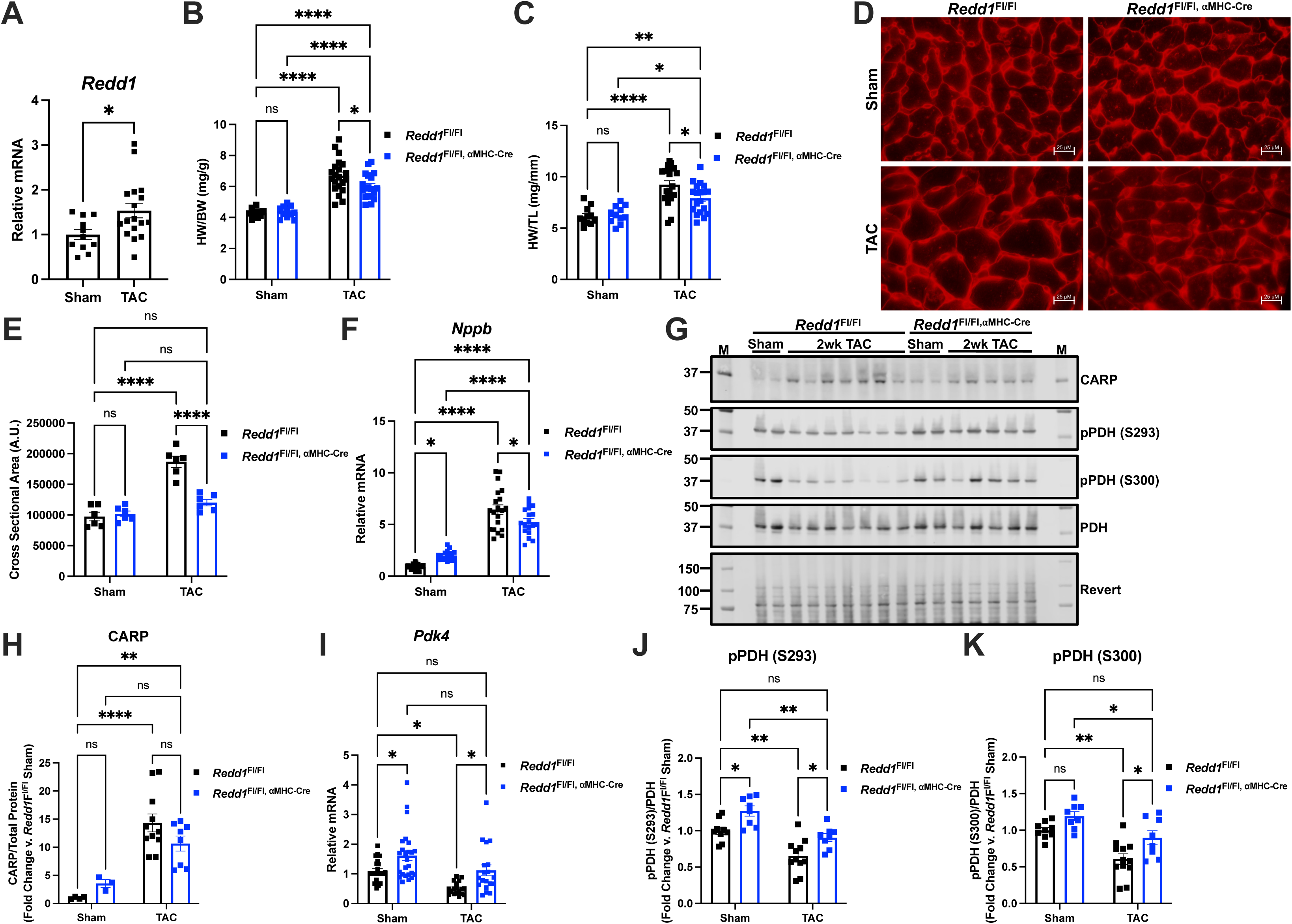
Cardiac REDD1 is required for metabolic and structural remodeling in pathological hypertrophy. Adult (10-12-week-old) male and female mice of the indicated genotypes were subjected to 2 weeks TAC or Sham operation. Hearts were then harvested. **A.** Total RNA extraction and qPCR for *Redd1* was performed (n=11,17), unpaired *t* test. **B-C.** Heart weights (HW) were normalized to **(B)** body weight (BW) or **(C)** tibia length (TL) and plotted as a ratio (mg/g or mg/mm, respectively). n=11,11,21,19 (HW/BW), n=11,11,21,19 (HW/TL), 2-way ANOVA. **D-E.** Hearts were fixed and stained with 594 wheat germ agglutinin. **(D)** Representative images are shown with 25 µm scale bars and **(E)** cardiomyocyte average cross-sectional area was measured. n=6,6,6,6, 2-way ANOVA. **F&I.** Total RNA extraction and qPCR for *Nppb* and *Pdk4* were performed. n=27,17,20,18 (*Nppb*), n=23,25,16,19 (*Pdk4*), 2-way ANOVA. **G-H&J-K.** Hearts were lysed and subjected to western blotting with the indicated antibodies. Signals were quantified with densitometry, normalized to **(H)** total protein or **(J-K)** total PDH, and plotted. n=4,3,11,8 (CARP), n=9,8,11,8 (pPDH (S293)), n=9,8,12,8 (pPDH (S300)), 2-way ANOVA. Error bars represent SEM. *p<0.05, **p<0.01, ****p<0.0001. M = marker.

Since the metabolic shift to enhanced glucose oxidation that occurs in the early adaptive phase of hypertrophic growth is supported by reduced PDK4^24^, we examined this metabolic remodeling in our *Redd1*^Fl/Fl^ or *Redd1*^Fl/Fl,αMHC-Cre^ mice subjected to Sham or TAC surgery. Indeed, we observed reduced *Pdk4* (**Fig 7I**), pPDH (S293) (**Fig 7G&J**), and pPDH (S300) (**Fig 7G&K**) levels in the hearts of our *Redd1*^Fl/Fl^ control mice subjected to TAC versus sham operation. Further, we observe significantly increased *Pdk4* (**Fig 7I**), pPDH (S293) (**Fig 7G&J**), and pPDH (S300) (**Fig 7G&K**) levels in the hearts of our *Redd1*^Fl/Fl,αMHC-Cre^ mice subjected to TAC versus *Redd1*^Fl/Fl^ control mice also subjected to TAC. Importantly, these levels were returned to those of control mice (*Redd1*^Fl/Fl^ sham-operated mice). Taken together, these findings suggest that pressure overload-induced REDD1 contributes’ to cardiac hypertrophy via its role in enhancing glucose oxidation.

## Discussion

Taken together, our findings outline a novel pathway whereby glucose- or pressure overload-induced REDD1 elevates glucose, while suppressing fatty acid metabolism (**Fig 8**). First, we show that glucose induces REDD1 (**Supp Fig 1**) and that, in this context, REDD1 suppresses PDK4 expression and PDH phosphorylation, as well as enhances PDH activity, effects which are required for glucose oxidation^19–21^ (**Fig 1A-H**, **Fig 2**). This is also evidenced by REDD1-mediated suppression of ECAR, which is an indirect measure of glycolysis and glycolytic capacity^50^, and elevation of MRC, which has been shown to positively correlate with glucose oxidation^39^ (**Fig 1I-M**). In addition, we demonstrate that REDD1 suppresses the expression of genes involved in fatty acid catabolism (**Fig 4**). Thus, we uniquely present REDD1 as a critical component of the glucose-induced switch in cardiac fuel metabolism, enhancing glucose oxidation and suppressing fatty acid catabolism.

**Figure 8.**
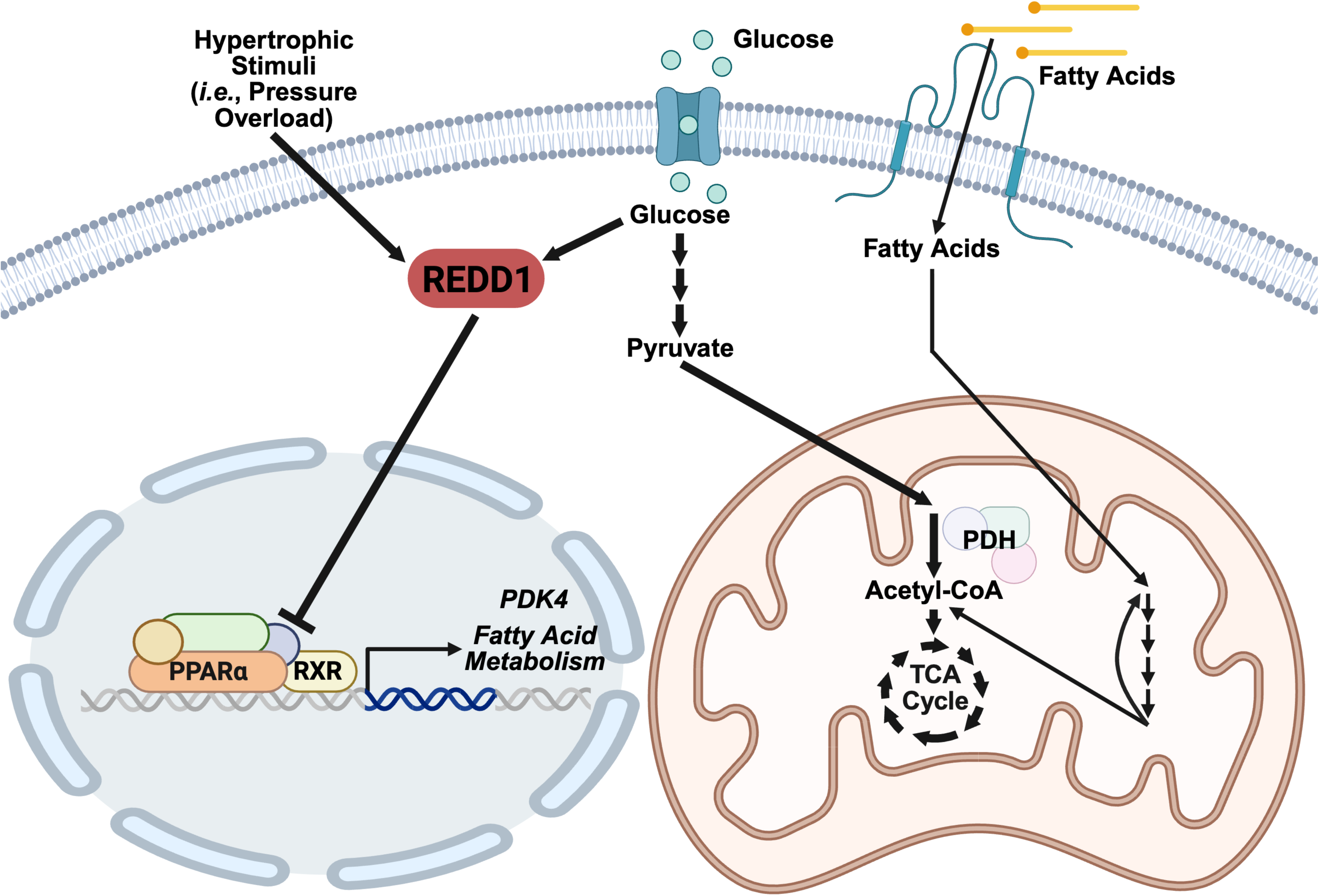
Summary. Our findings outline a mechanism whereby pressure overload- or glucose-induced REDD1 is critical for activating glucose and suppressing fatty acid oxidation pathways. Specifically, elevated REDD1 inhibits PPARα, thus inhibiting the expression of *PDK4* and fatty acid catabolic genes. We also show that this is independent of REDD1’s ability to inhibit mTORC1.

In addition to uncovering a role for REDD1 in increasing the ratio of glucose-to-fatty acid fuel use in the heart, we also outline the mechanisms by which this occurs. Specifically, we demonstrate that REDD1’s inhibition of PPARα is at least partially responsible for the suppression of *PDK4* and fatty acid catabolic genes (**Fig 6**). Additionally, we demonstrate that these effects are not dependent on REDD1’s ability to inhibit mTORC1 (**Fig 3&5A-D**), which is critical because this is the established function of REDD1^11^. It is important to note that the mechanisms by which REDD1 inhibits PPARα remain unclear. It has been shown that REDD1 is present in the nuclei of cells^51^, but its function here is not well studied. It is possible, however, that glucose-inducible REDD1 is nuclear and directly binds PPARα, inhibiting its transcriptional activity.

Beyond elucidating a role for REDD1 in cardiac physiology and establishing its underlying mechanism, we examined REDD1 in a model of cardiac pathology. We show that pressure overload, as is observed in hypertension, induces cardiac REDD1 expression (**Fig 7A**) and that this contributes to pathological cardiac hypertrophy (**Fig 7B-H**). It is well established that a metabolic shift, characterized by enhanced glucose and suppressed fatty acid oxidation, is required for early adaptive hypertrophic growth in response to pressure overload^24–27^. It is also evident that reduced PPARα is critical for reduced expression of PDK4 and fatty acid catabolic genes, therefore contributing to elevated glucose and suppressed fatty acid oxidation in this context, respectively. Herein, we demonstrate that REDD1 is required for pressure overload-induced reductions in PDK4 and PDH phosphorylation, which are critical for glucose oxidation^19–21^(**Fig 7G&I-K**). Thus, these findings reinforce a role for REDD1 as major metabolic switch. Further, these data are consistent with an mTORC1-independent role for REDD1 in cardiac hypertrophy, as elevated mTORC1 signaling, as observed with REDD1 deletion, exacerbates hypertrophic growth^52, 53^. Importantly, the mechanisms of REDD1 induction in hypertrophy, as well as the role of PPARα in the associated gene expression changes, remain uninvestigated. Finally, our findings position REDD1 as a potential therapeutic target for pathological cardiac remodeling and failure. While REDD1 inhibition in the early adaptive phase of hypertrophic growth may be ill-advised, examining its levels in early heart failure may allow for its inhibition in this context, as PPARα activation in this period has been shown to preserve cardiac energetics and function^54^.

The Randle Cycle describes a phenomenon in which glucose and fatty acids compete as metabolic substrates in a given tissue^55^. For example, when glucose utilization is high, there is an accumulation of cytosolic malonyl CoA, which is derived from citrate and acetyl CoA, and inhibits mitochondrial fatty acid uptake via carnitine palmitoyltransferase 1 (CPT1)^56^. Conversely, fatty acid oxidation results in the accumulation of cytosolic citrate that inhibits phosphofructokinase activity and, therefore, glycolysis^56^. PPARα is activated by fatty acids^45^ and increases the expression of target gene, *PDK4*^17, 18, 22^, which inhibits glucose oxidation and is, therefore, also an established component of the Randle Cycle. Since we outline a role for REDD1 as a glucose-inducible endogenous PPARα inhibitor, we present REDD1 as a novel component of the Randle Cycle.

While our work examines the role of REDD1 in cardiac metabolism, it is likely that REDD1 also controls fuel use in other tissues. While both REDD1 and PPARα are ubiquitously expressed, basal metabolic fuel preference, metabolic flexibility, and substrate storage capacity for a particular tissue may alter the functional relevance of this pathway. For example, glucose-induced REDD1 and its ability to promote glucose oxidation and suppress fatty acid catabolism via PPARα inhibition likely also occurs in the liver and skeletal muscle. In these tissues and in contrast to the heart, this pathway is also likely to suppress glycogen synthesis while promoting glycogenolysis. In a metabolically inflexible tissue that has a high reliance on glucose, such as the brain^57^, we expect that this pathway is highly relevant at baseline, supporting glucose oxidation.

In addition to tissue-specific relevance, the outlined pathway is also expected to be critical in the context of various physiological stimuli and pathological stressors. For example, REDD1 has been shown to be induced by insulin^5, 6^ and exercise^7^ in numerous tissues. It is possible that insulin-induced REDD1 also enhances glucose oxidation and suppresses fatty acid catabolism via this PPARα-dependent mechanism. Notably, REDD1 is also upregulated in type II diabetes^9, 58^, however, the applicability of this pathway in this context remains unexplored. Interestingly, similar to pathological (*i.e.*, pressure overload-induced)^3, 59^ cardiac hypertrophic growth, physiological (*i.e.*, exercise-induced)^60^ cardiac hypertrophy is also characterized by a switch from fatty acid to glucose utilization. Exercise-induced REDD1 upregulation^61^ likely also contributes to this shift in fuel preference. Finally, several studies report upregulation, while others demonstrate downregulation of REDD1 in response to hypoxia or ischemia^62–64^. It will be interesting to examine REDD1 in this context and determine if PPARα inhibition plays a role.

Overall, we present REDD1 as a critical regulator of cardiac metabolic fuel preference, whereby it enhances glucose oxidation while reducing fatty acid catabolism, via its inhibition of PPARα. We also demonstrate that this occurs independent of REDD1-mediated mTORC1 inhibition. Finally, we show that REDD1 is a major contributor to the metabolic and structural remodeling that occurs in pathological cardiac hypertrophy. Moving forward, more work is required to uncover the mechanisms by which REDD1 inhibits PPARα activity, as well as the role of this pathway in diverse tissues during various physiological conditions or during pathology. These studies are critical for understanding metabolic fuel preference and targeting it for therapeutic benefit.

## Acknowledgements

The authors thank Dr. David L. Williamson, PhD (Pennsylvania State University at Harrisburg) for the *Redd1*^Fl/Fl^ mice, as well as Dr. Alexey Zaitsev, PhD and Steven Burrows (FBRI at Virginia Tech Carilion) for their contributions to the metabolomics processing and analysis. BioRender was used for the generation of Figure 8.

## References

1. Lopaschuk GD, Karwi QG, Tian R, Wende AR, Abel ED. Cardiac Energy Metabolism in Heart Failure. Circulation Research. 2021;128(10):1487–513. doi: 10.1161/CIRCRESAHA.121.318241.

2. Bertrand L, Horman S, Beauloye C, Vanoverschelde J-L. Insulin signalling in the heart. Cardiovascular Research. 2008;79(2):238–48. doi: 10.1093/cvr/cvn093.

3. Allard MF, Schonekess BO, Henning SL, English DR, Lopaschuk GD. Contribution of oxidative metabolism and glycolysis to ATP production in hypertrophied hearts. American Journal of Physiology-Heart and Circulatory Physiology. 1994;267(2):H742–H50. doi: 10.1152/ajpheart.1994.267.2.H742. PubMed PMID: 8067430.

4. Stevens SA, Gonzalez Aguiar MK, Toro AL, Yerlikaya EI, Sunilkumar S, VanCleave AM, Pfleger J, Bradley EA, Kimball SR, Dennis MD. PERK/ATF4-dependent expression of the stress response protein REDD1 promotes proinflammatory cytokine expression in the heart of obese mice. American Journal of Physiology-Endocrinology and Metabolism. 2023;324(1):E62–E72. doi: 10.1152/ajpendo.00238.2022. PubMed PMID: 36383638.

5. Regazzetti C, Bost F, Le Marchand-Brustel Y, Tanti J-F, Giorgetti-Peraldi S. Insulin Induces REDD1 Expression through Hypoxia-inducible Factor 1 Activation in Adipocytes *. Journal of Biological Chemistry. 2010;285(8):5157–64. doi: 10.1074/jbc.M109.047688.

6. Regazzetti C, Dumas K, Le Marchand-Brustel Y, Peraldi P, Tanti J-F, Giorgetti-Peraldi S. Regulated in Development and DNA Damage Responses -1 (REDD1) Protein Contributes to Insulin Signaling Pathway in Adipocytes. PLOS ONE. 2012;7(12):e52154. doi: 10.1371/journal.pone.0052154.

7. Gordon BS, Steiner JL, Rossetti ML, Qiao S, Ellisen LW, Govindarajan SS, Eroshkin AM, Williamson DL, Coen PM. REDD1 induction regulates the skeletal muscle gene expression signature following acute aerobic exercise. American Journal of Physiology-Endocrinology and Metabolism. 2017;313(6):E737–E47. doi: 10.1152/ajpendo.00120.2017. PubMed PMID: 28899858.

8. Hulmi JJ, Silvennoinen M, Lehti M, Kivelä R, Kainulainen H. Altered REDD1, myostatin, and Akt/mTOR/FoxO/MAPK signaling in streptozotocin-induced diabetic muscle atrophy. American Journal of Physiology-Endocrinology and Metabolism. 2012;302(3):E307–E15. doi: 10.1152/ajpendo.00398.2011. PubMed PMID: 22068602.

9. Williamson DL, Li Z, Tuder RM, Feinstein E, Kimball SR, Dungan CM. Altered nutrient response of mTORC1 as a result of changes in REDD1 expression: effect of obesity vs. REDD1 deficiency. J Appl Physiol (1985). 2014;117(3):246–56. Epub 20140529. doi: 10.1152/japplphysiol.01350.2013. PubMed PMID: 24876363; PMCID: PMC4122690.

10. Choi HS, Ahn JH, Park JH, Won MH, Lee CH. Age-dependent changes in the protein expression levels of Redd1 and mTOR in the gerbil hippocampus during normal aging. Mol Med Rep. 2016;13(3):2409–14. doi: 10.3892/mmr.2016.4835.

11. Brugarolas J, Lei K, Hurley RL, Manning BD, Reiling JH, Hafen E, Witters LA, Ellisen LW, Kaelin WG. Regulation of mTOR function in response to hypoxia by REDD1 and the TSC1/TSC2 tumor suppressor complex. Genes & Development. 2004;18(23):2893–904. doi: 10.1101/gad.1256804.

12. Laplante M, Sabatini DM. mTOR signaling at a glance. J Cell Sci. 2009;122(Pt 20):3589–94. doi: 10.1242/jcs.051011. PubMed PMID: 19812304; PMCID: PMC2758797.

13. Dungan CM, Wright DC, Williamson DL. Lack of REDD1 reduces whole body glucose and insulin tolerance, and impairs skeletal muscle insulin signaling. Biochemical and Biophysical Research Communications. 2014;453(4):778–83. doi: 10.1016/j.bbrc.2014.10.032.

14. Dennis MD, Kimball SR, Fort PE, Jefferson LS. Regulated in Development and DNA Damage 1 Is Necessary for Hyperglycemia-induced Vascular Endothelial Growth Factor Expression in the Retina of Diabetic Rodents. Journal of Biological Chemistry. 2015;290(6):3865–74. doi: 10.1074/jbc.M114.623058.

15. Sunilkumar S, Toro AL, McCurry CM, VanCleave AM, Stevens SA, Miller WP, Kimball SR, Dennis MD. Stress response protein REDD1 promotes diabetes-induced retinal inflammation by sustaining canonical NF-&#x3ba;B signaling. Journal of Biological Chemistry. 2022;298(12). doi: 10.1016/j.jbc.2022.102638.

16. Zhu Y, Soto J, Anderson B, Riehle C, Zhang YC, Wende AR, Jones D, McClain DA, Abel ED. Regulation of fatty acid metabolism by mTOR in adult murine hearts occurs independently of changes in PGC-1α. American Journal of Physiology-Heart and Circulatory Physiology. 2013;305(1):H41–H51. doi: 10.1152/ajpheart.00877.2012. PubMed PMID: 23624629.

17. Rakhshandehroo M, Sanderson LM, Matilainen M, Stienstra R, Carlberg C, de Groot PJ, Muller M, Kersten S. Comprehensive analysis of PPARalpha-dependent regulation of hepatic lipid metabolism by expression profiling. PPAR Res. 2007;2007:26839. doi: 10.1155/2007/26839. PubMed PMID: 18288265; PMCID: PMC2233741.

18. Wu P, Peters JM, Harris RA. Adaptive Increase in Pyruvate Dehydrogenase Kinase 4 during Starvation Is Mediated by Peroxisome Proliferator-Activated Receptor α. Biochemical and Biophysical Research Communications. 2001;287(2):391–6. doi: 10.1006/bbrc.2001.5608.

19. Gopal K, Almutairi M, Al Batran R, Eaton F, Gandhi M, Ussher JR. Cardiac-Specific Deletion of Pyruvate Dehydrogenase Impairs Glucose Oxidation Rates and Induces Diastolic Dysfunction. Front Cardiovasc Med. 2018;5:17. Epub 20180306. doi: 10.3389/fcvm.2018.00017. PubMed PMID: 29560354; PMCID: PMC5845646.

20. Linn TC, Pettit FH, Reed LJ. a-Keto Acid Dehydrogenase Complexes, X. Regulation of the Activity of the Pyruvate Dehydrogenase Complex from Beef Kidney Mitochondria by Phosphorylation and Dephosphorylation. Proceedings of the National Academy of Sciences. 1969;62(1):234–41. doi: doi:10.1073/pnas.62.1.234.

21. Patel MS, Korotchkina LG. Regulation of mammalian pyruvate dehydrogenase complex by phosphorylation: complexity of multiple phosphorylation sites and kinases. Experimental & Molecular Medicine. 2001;33(4):191–7. doi: 10.1038/emm.2001.32.

22. Rakhshandehroo M, Knoch B, Müller M, Kersten S. Peroxisome proliferator-activated receptor alpha target genes. PPAR Res. 2010;2010. Epub 20100926. doi: 10.1155/2010/612089. PubMed PMID: 20936127; PMCID: PMC2948931.

23. van der Meer DLM, Degenhardt T, Väisänen S, de Groot PJ, Heinäniemi M, de Vries SC, Müller M, Carlberg C, Kersten S. Profiling of promoter occupancy by PPARα in human hepatoma cells via ChIP-chip analysis. Nucleic Acids Research. 2010;38(9):2839–50. doi: 10.1093/nar/gkq012.

24. Akki A, Smith K, Seymour AM. Compensated cardiac hypertrophy is characterised by a decline in palmitate oxidation. Mol Cell Biochem. 2008;311(1-2):215–24. Epub 20080216. doi: 10.1007/s11010-008-9711-y. PubMed PMID: 18278440.

25. Christe ME, Rodgers RL. Altered glucose and fatty acid oxidation in hearts of the spontaneously hypertensive rat. J Mol Cell Cardiol. 1994;26(10):1371–5. doi: 10.1006/jmcc.1994.1155. PubMed PMID: 7869397.

26. Doenst T, Pytel G, Schrepper A, Amorim P, Farber G, Shingu Y, Mohr FW, Schwarzer M. Decreased rates of substrate oxidation ex vivo predict the onset of heart failure and contractile dysfunction in rats with pressure overload. Cardiovasc Res. 2010;86(3):461–70. Epub 20091224. doi: 10.1093/cvr/cvp414. PubMed PMID: 20035032.

27. Kolwicz SC, Jr., Olson DP, Marney LC, Garcia-Menendez L, Synovec RE, Tian R. Cardiac-specific deletion of acetyl CoA carboxylase 2 prevents metabolic remodeling during pressure-overload hypertrophy. Circ Res. 2012;111(6):728–38. Epub 20120622. doi: 10.1161/CIRCRESAHA.112.268128. PubMed PMID: 22730442; PMCID: PMC3434870.

28. Dai W, Miller WP, Toro AL, Black AJ, Dierschke SK, Feehan RP, Kimball SR, Dennis MD. Deletion of the stress-response protein REDD1 promotes ceramide-induced retinal cell death and JNK activation. Faseb j. 2018;32(12):fj201800413RR. Epub 20180619. doi: 10.1096/fj.201800413RR. PubMed PMID: 29920218; PMCID: PMC6219834.

29. Ackers-Johnson M, Li PY, Holmes AP, O’Brien S-M, Pavlovic D, Foo RS. A Simplified, Langendorff-Free Method for Concomitant Isolation of Viable Cardiac Myocytes and Nonmyocytes From the Adult Mouse Heart. Circulation Research. 2016;119(8):909–20. doi: doi:10.1161/CIRCRESAHA.116.309202.

30. Brafman A, Mett I, Shafir M, Gottlieb H, Damari G, Gozlan-Kelner S, Vishnevskia-Dai V, Skaliter R, Einat P, Faerman A, Feinstein E, Shoshani T. Inhibition of Oxygen-Induced Retinopathy in RTP801-Deficient Mice. Investigative Ophthalmology & Visual Science. 2004;45(10):3796–805. doi: 10.1167/iovs.04-0052.

31. Pereira RO, Wende AR, Crum A, Hunter D, Olsen CD, Rawlings T, Riehle C, Ward WF, Abel ED. Maintaining PGC-1alpha expression following pressure overload-induced cardiac hypertrophy preserves angiogenesis but not contractile or mitochondrial function. FASEB J. 2014;28(8):3691–702. Epub 20140428. doi: 10.1096/fj.14-253823. PubMed PMID: 24776744; PMCID: PMC4101649.

32. Hem A, Smith AJ, Solberg P. Saphenous vein puncture for blood sampling of the mouse, rat, hamster, gerbil, guineapig, ferret and mink. Laboratory Animals. 1998;32(4):364–8. doi: 10.1258/002367798780599866. PubMed PMID: 9807749.

33. Oka SI, Sreedevi K, Shankar TS, Yedla S, Arowa S, James A, Stone KG, Olmos K, Sabry AD, Horiuchi A, Cawley KM, O’Very S A, Tong M, Byun J, Xu X, Kashyap S, Mourad Y, Vehra O, Calder D, Lunde T, Liu T, Li H, Mashchek JA, Cox J, Saijoh Y, Drakos SG, Warren JS. PERM1 regulates energy metabolism in the heart via ERRalpha/PGC-1alpha axis. Front Cardiovasc Med. 2022;9:1033457. Epub 20221107. doi: 10.3389/fcvm.2022.1033457. PubMed PMID: 36419485; PMCID: PMC9676655.

34. Shibayama J, Taylor TG, Venable PW, Rhodes NL, Gil RB, Warren M, Wende AR, Abel ED, Cox J, Spitzer KW, Zaitsev AV. Metabolic determinants of electrical failure in ex-vivo canine model of cardiac arrest: evidence for the protective role of inorganic pyrophosphate. PLoS One. 2013;8(3):e57821. Epub 20130308. doi: 10.1371/journal.pone.0057821. PubMed PMID: 23520482; PMCID: PMC3592894.

35. Shibayama J, Yuzyuk TN, Cox J, Makaju A, Miller M, Lichter J, Li H, Leavy JD, Franklin S, Zaitsev AV. Metabolic remodeling in moderate synchronous versus dyssynchronous pacing-induced heart failure: integrated metabolomics and proteomics study. PLoS One. 2015;10(3):e0118974. Epub 20150319. doi: 10.1371/journal.pone.0118974. PubMed PMID: 25790351; PMCID: PMC4366225.

36. Warren JS, Tracy CM, Miller MR, Makaju A, Szulik MW, Oka SI, Yuzyuk TN, Cox JE, Kumar A, Lozier BK, Wang L, Llana JG, Sabry AD, Cawley KM, Barton DW, Han YH, Boudina S, Fiehn O, Tucker HO, Zaitsev AV, Franklin S. Histone methyltransferase Smyd1 regulates mitochondrial energetics in the heart. Proc Natl Acad Sci U S A. 2018;115(33):E7871–E80. Epub 20180730. doi: 10.1073/pnas.1800680115. PubMed PMID: 30061404; PMCID: PMC6099878.

37. Rabinowitz JD, Enerbäck S. Lactate: the ugly duckling of energy metabolism. Nature Metabolism. 2020;2(7):566–71. doi: 10.1038/s42255-020-0243-4.

38. Benjamin D, Robay D, Hindupur SK, Pohlmann J, Colombi M, El-Shemerly MY, Maira S-M, Moroni C, Lane HA, Hall MN. Dual Inhibition of the Lactate Transporters MCT1 and MCT4 Is Synthetic Lethal with Metformin due to NAD+ Depletion in Cancer Cells. Cell Reports. 2018;25(11):3047–58.e4. doi: 10.1016/j.celrep.2018.11.043.

39. Pfleger J, He M, Abdellatif M. Mitochondrial complex II is a source of the reserve respiratory capacity that is regulated by metabolic sensors and promotes cell survival. Cell Death Dis. 2015;6(7):e1835. Epub 20150730. doi: 10.1038/cddis.2015.202. PubMed PMID: 26225774; PMCID: PMC4650745.

40. Shoshani T, Alexander F, Igor M, Elena Z, Tamar T, Svetlana G, Yana M, Shlomo E, Andrei B, Ayelet C, Hagar K, Iris K, Ada R, Orna M, Eli K, Dena L, Paz E, Rami S, and Feinstein E. Identification of a Novel Hypoxia-Inducible Factor 1-Responsive Gene, RTP801, Involved in Apoptosis. Molecular and Cellular Biology. 2002;22(7):2283–93. doi: 10.1128/MCB.22.7.2283-2293.2002.

41. Stevens SA, Sunilkumar S, Subrahmanian SM, Toro AL, Cavus O, Omorogbe EV, Bradley EA, Dennis MD. REDD1 Deletion Suppresses NF-κB Signaling in Cardiomyocytes and Prevents Deficits in Cardiac Function in Diabetic Mice. Int J Mol Sci. 2024;25(12). Epub 20240612. doi: 10.3390/ijms25126461. PubMed PMID: 38928166; PMCID: PMC11204184.

42. Arriola Apelo SI, Neuman JC, Baar EL, Syed FA, Cummings NE, Brar HK, Pumper CP, Kimple ME, Lamming DW. Alternative rapamycin treatment regimens mitigate the impact of rapamycin on glucose homeostasis and the immune system. Aging Cell. 2016;15(1):28–38. Epub 20151013. doi: 10.1111/acel.12405. PubMed PMID: 26463117; PMCID: PMC4717280.

43. Tan CY, Hagen T. mTORC1 dependent regulation of REDD1 protein stability. PLoS One. 2013;8(5):e63970. Epub 20130522. doi: 10.1371/journal.pone.0063970. PubMed PMID: 23717519; PMCID: PMC3661664.

44. Sengupta S, Peterson TR, Laplante M, Oh S, Sabatini DM. mTORC1 controls fasting-induced ketogenesis and its modulation by ageing. Nature. 2010;468(7327):1100–4. doi: 10.1038/nature09584.

45. Liu J, Sahin C, Ahmad S, Magomedova L, Zhang M, Jia Z, Metherel AH, Orellana A, Poda G, Bazinet RP, Attisano L, Cummins CL, Peng H, Krause HM. The omega-3 hydroxy fatty acid 7(*S)*-HDHA is a high-affinity PPARa ligand that regulates brain neuronal morphology. Science Signaling. 2022;15(741):eabo1857. doi: doi:10.1126/scisignal.abo1857.

46. Walker MA, Chen H, Yadav A, Ritterhoff J, Villet O, McMillen T, Wang Y, Purcell H, Djukovic D, Raftery D, Isoherranen N, Tian R. Raising NAD^+^ Level Stimulates Short-Chain Dehydrogenase/Reductase Proteins to Alleviate Heart Failure Independent of Mitochondrial Protein Deacetylation. Circulation. 2023;148(25):2038–57. doi: doi:10.1161/CIRCULATIONAHA.123.066039.

47. Xu HE, Stanley TB, Montana VG, Lambert MH, Shearer BG, Cobb JE, McKee DD, Galardi CM, Plunket KD, Nolte RT, Parks DJ, Moore JT, Kliewer SA, Willson TM, Stimmel JB. Structural basis for antagonist-mediated recruitment of nuclear co-repressors by PPARα. Nature. 2002;415(6873):813–7. doi: 10.1038/415813a.

48. Sayed D, Yang Z, He M, Pfleger JM, Abdellatif M. Acute targeting of general transcription factor IIB restricts cardiac hypertrophy via selective inhibition of gene transcription. Circ Heart Fail. 2015;8(1):138–48. Epub 20141114. doi: 10.1161/CIRCHEARTFAILURE.114.001660. PubMed PMID: 25398966; PMCID: PMC4401077.

49. Chen Y, Guo H, Xu D, Xu X, Wang H, Hu X, Lu Z, Kwak D, Xu Y, Gunther R, Huo Y, Weir EK. Left ventricular failure produces profound lung remodeling and pulmonary hypertension in mice: heart failure causes severe lung disease. Hypertension. 2012;59(6):1170–8. Epub 20120416. doi: 10.1161/HYPERTENSIONAHA.111.186072. PubMed PMID: 22508832; PMCID: PMC3402091.

50. Divakaruni AS, Paradyse A, Ferrick DA, Murphy AN, Jastroch M. Chapter Sixteen - Analysis and Interpretation of Microplate-Based Oxygen Consumption and pH Data. In: Murphy AN, Chan DC, editors. Methods in Enzymology: Academic Press; 2014. p. 309–54.

51. Lin L, Stringfield TM, Shi X, Chen Y. Arsenite induces a cell stress-response gene, RTP801, through reactive oxygen species and transcription factors Elk-1 and CCAAT/enhancer-binding protein. Biochem J. 2005;392(Pt 1):93–102. doi: 10.1042/BJ20050553. PubMed PMID: 16008523; PMCID: PMC1317668.

52. Shende P, Plaisance I, Morandi C, Pellieux C, Berthonneche C, Zorzato F, Krishnan J, Lerch R, Hall MN, Ruegg MA, Pedrazzini T, Brink M. Cardiac raptor ablation impairs adaptive hypertrophy, alters metabolic gene expression, and causes heart failure in mice. Circulation. 2011;123(10):1073–82. Epub 20110228. doi: 10.1161/CIRCULATIONAHA.110.977066. PubMed PMID: 21357822.

53. Zhang D, Contu R, Latronico MV, Zhang J, Rizzi R, Catalucci D, Miyamoto S, Huang K, Ceci M, Gu Y, Dalton ND, Peterson KL, Guan KL, Brown JH, Chen J, Sonenberg N, Condorelli G. MTORC1 regulates cardiac function and myocyte survival through 4E-BP1 inhibition in mice. J Clin Invest. 2010;120(8):2805–16. Epub 20100719. doi: 10.1172/JCI43008. PubMed PMID: 20644257; PMCID: PMC2912201.

54. Kaimoto S, Hoshino A, Ariyoshi M, Okawa Y, Tateishi S, Ono K, Uchihashi M, Fukai K, Iwai-Kanai E, Matoba S. Activation of PPAR-alpha in the early stage of heart failure maintained myocardial function and energetics in pressure-overload heart failure. Am J Physiol Heart Circ Physiol. 2017;312(2):H305–H13. Epub 20161223. doi: 10.1152/ajpheart.00553.2016. PubMed PMID: 28011586.

55. Randle PJ, Garland PB, Hales CN, Newsholme EA. The Glucose Fatty-Acid Cycle its Role in Insulin Sensitivity and the Metabolic Disturbances of Diabetes Mellitus. The Lancet. 1963;281(7285):785–9. doi: 10.1016/S0140-6736(63)91500-9.

56. Hue L, Taegtmeyer H. The Randle cycle revisited: a new head for an old hat. Am J Physiol Endocrinol Metab. 2009;297(3):E578–91. Epub 20090616. doi: 10.1152/ajpendo.00093.2009. PubMed PMID: 19531645; PMCID: PMC2739696.

57. Himwich H, Nahum L. The respiratory quotient of the brain. American Journal of Physiology-Legacy Content. 1932;101(3):446–53.

58. Williamson DL, Dungan CM, Mahmoud AM, Mey JT, Blackburn BK, Haus JM. Aberrant REDD1-mTORC1 responses to insulin in skeletal muscle from Type 2 diabetics. American Journal of Physiology-Regulatory, Integrative and Comparative Physiology. 2015;309(8):R855–R63. doi: 10.1152/ajpregu.00285.2015. PubMed PMID: 26269521.

59. Akki A, Smith K, Seymour A-ML. Compensated cardiac hypertrophy is characterised by a decline in palmitate oxidation. Molecular and Cellular Biochemistry. 2008;311(1):215–24. doi: 10.1007/s11010-008-9711-y.

60. Gertz EW, Wisneski JA, Stanley WC, Neese RA. Myocardial substrate utilization during exercise in humans. Dual carbon-labeled carbohydrate isotope experiments. The Journal of Clinical Investigation. 1988;82(6):2017–25. doi: 10.1172/JCI113822.

61. Gordon BS, Steiner JL, Rossetti ML, Qiao S, Ellisen LW, Govindarajan SS, Eroshkin AM, Williamson DL, Coen PM. REDD1 induction regulates the skeletal muscle gene expression signature following acute aerobic exercise. Am J Physiol Endocrinol Metab. 2017;313(6):E737–E47. Epub 20170912. doi: 10.1152/ajpendo.00120.2017. PubMed PMID: 28899858; PMCID: PMC5814598.

62. Seong M, Lee J, Kang H. Hypoxia-induced regulation of mTOR signaling by miR-7 targeting REDD1. Journal of Cellular Biochemistry. 2019;120(3):4523–32. doi: 10.1002/jcb.27740.

63. Schwarzer R, Tondera D, Arnold W, Giese K, Klippel A, Kaufmann J. REDD1 integrates hypoxia-mediated survival signaling downstream of phosphatidylinositol 3-kinase. Oncogene. 2005;24(7):1138–49. doi: 10.1038/sj.onc.1208236.

64. Huang P, Fu J, Chen L, Ju C, Wu K, Liu H, Liu Y, Qi B, Qi B, Liu L. Redd1 protects against post-infarction cardiac dysfunction by targeting apoptosis and autophagy. Int J Mol Med. 2019;44(6):2065–76. Epub 20191004. doi: 10.3892/ijmm.2019.4366. PubMed PMID: 31638187; PMCID: PMC6844599.

