## Supplemental Data for "Cardiac REDD1 alters glucose and fatty acid metabolic gene expression via an mTORC1-independent, PPARα-dependent mechanism and drives hypertrophic growth"

Short Title: Cardiac REDD1 alters metabolic genes

Correspondence (Present Address): Jessica Pfleger

1 Baylor Plaza

Houston, TX 77030

Supplemental Material:

- Supplemental Table 1
- Supplemental Figures 1-5

### Supplemental Figure 1.

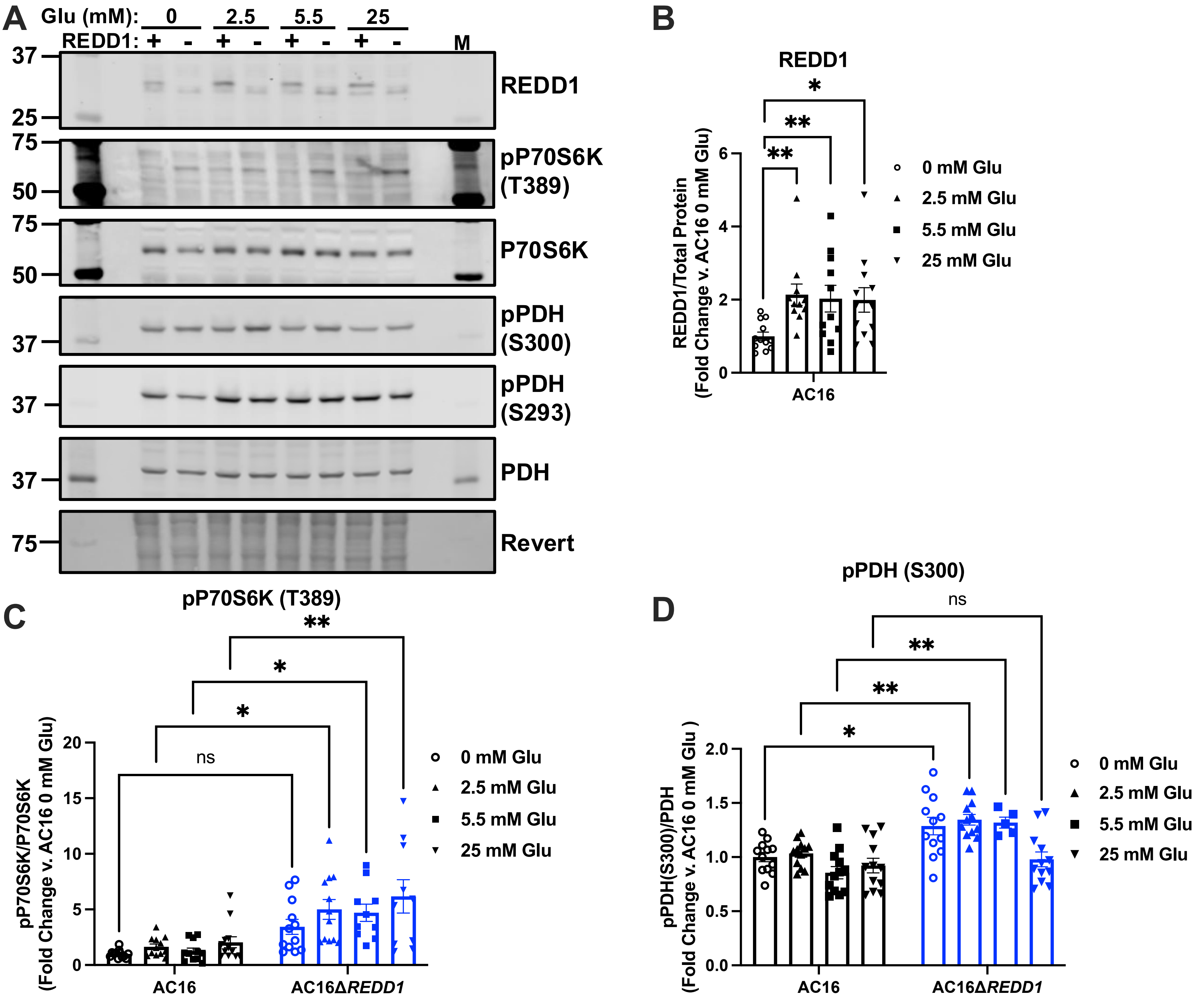

**Supplemental Figure 1. Cardiomyocyte REDD1 is induced by glucose, inhibits mTORC1, and suppresses PDH phosphorylation at physiological glucose levels. A-D.** AC16 and AC16Δ*REDD1* cardiomyocytes were cultured in DMEM, no glucose, un-supplemented (0 mM Glucose) or DMEM, no glucose supplemented with 2.5, 5.5, or 25 mM glucose for 24 hours and subjected to western blotting with the indicated antibodies. Signals were quantified with densitometry, normalized to total protein, P70S6K, or PDH as indicated and plotted. n=12,11,11,12 (REDD1), n=12,12,12,12,11,10,12,10 (pP70S6K (T389)), n=12,12,12,12,12,5,12,12 (pPDH (S300)) 1-way ANOVA, 2-way ANOVA. Error bars represent SEM. *p<0.05, **p<0.01. M = marker.

### Supplemental Figure 2.

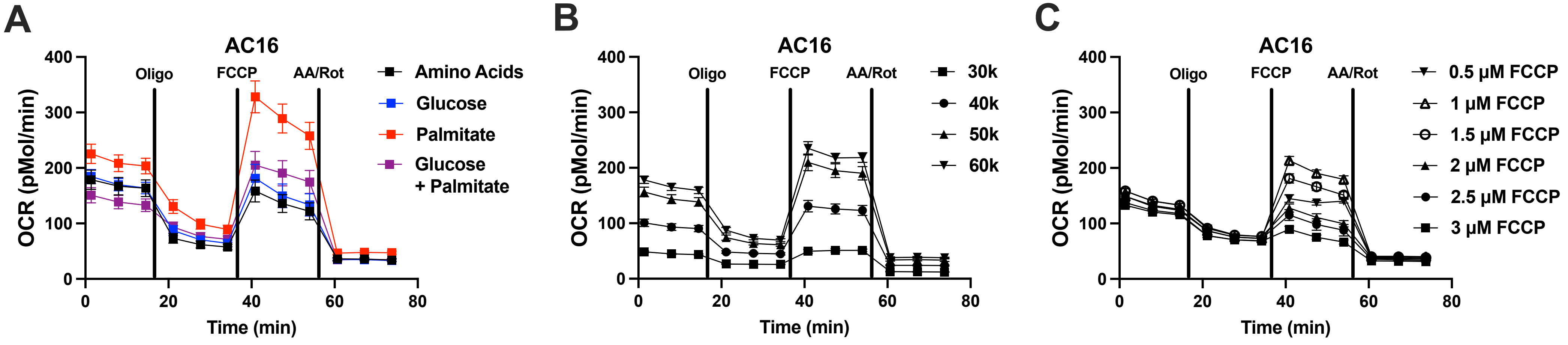

**Supplemental Figure 2. AC16 cardiomyocytes have an adult-like metabolic phenotype. A-C.** AC16 cardiomyocytes were plated at a density of **(A)** 60,000 cells per well and treated with the indicated substrates (DMEM, no glucose (amino acids) or DMEM, no glucose supplemented 5.5 mM glucose, 200 µM palmitate, or 5.5 mM glucose and 200 µM palmitate), **(B)** 30,000, 40,000, 50,000, or 60,000 cells per well, treated with glucose and palmitate, or **(C)** 60,000 cells per well treated with glucose and palmitate with the indicated dose of FCCP (0.5, 1, 1.5, 2, 2.5, or 3 µM), and subjected to mitochondrial stress tests as detailed in the methods section. The results are plotted as oxygen consumption (OCR) (pMol/min) versus time (minutes). n=22,21,22,20 **(A)**, n=10 **(B)**, and n=10,9,10,10,10,10 **(C)**.

### Supplemental Figure 3.

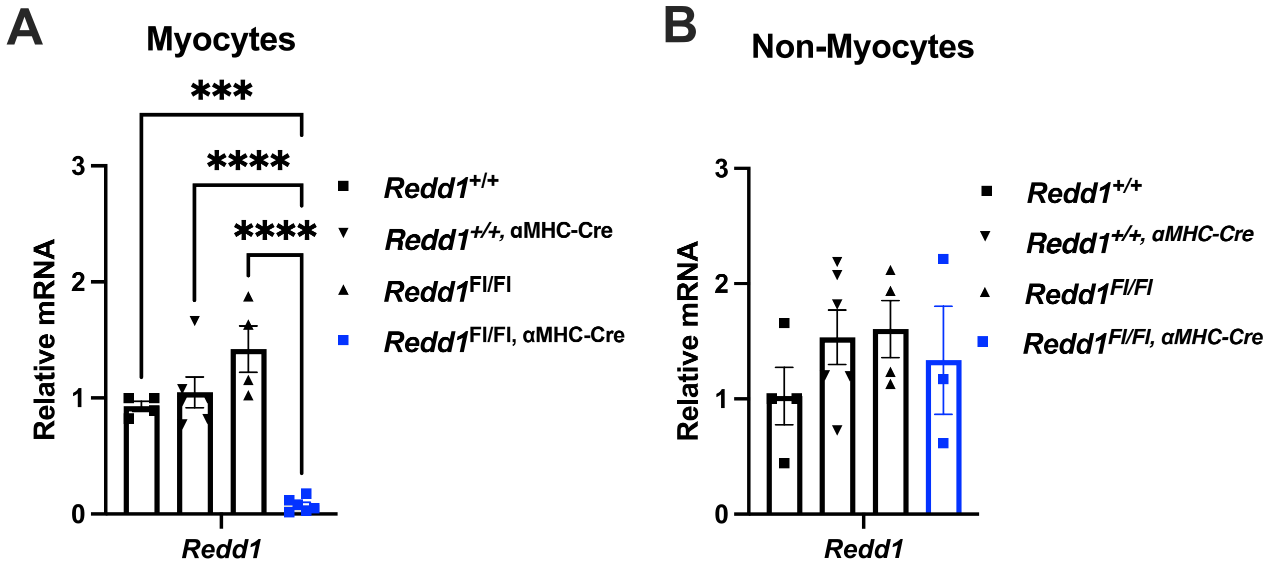

**Supplemental Figure 3. REDD1 is specifically deleted from cardiomyocytes *in vivo.* A-B.** Hearts of adult (8-14-week-old) male and female *Redd1*$^{+/+}$, *Redd1*$^{+/+, \alpha MHC\text{-}Cre}$, *Redd1*$^{Fl/Fl}$, and *Redd1*$^{Fl/Fl, \alpha MHC\text{-}Cre}$ mice were subjected to cardiomyocyte isolation as detailed in the methods section. The **(A)** myocyte and **(B)** non-myocyte fractions were subjected to total RNA extraction and qPCR for *Redd1*. n=4,6,4,6 (Myocyte), n=4,6,4,3 (Non-Myocyte), 1-way ANOVA. Error bars represent SEM. ***$p<0.001$, ****$p<0.0001$.

### Supplemental Figure 4.

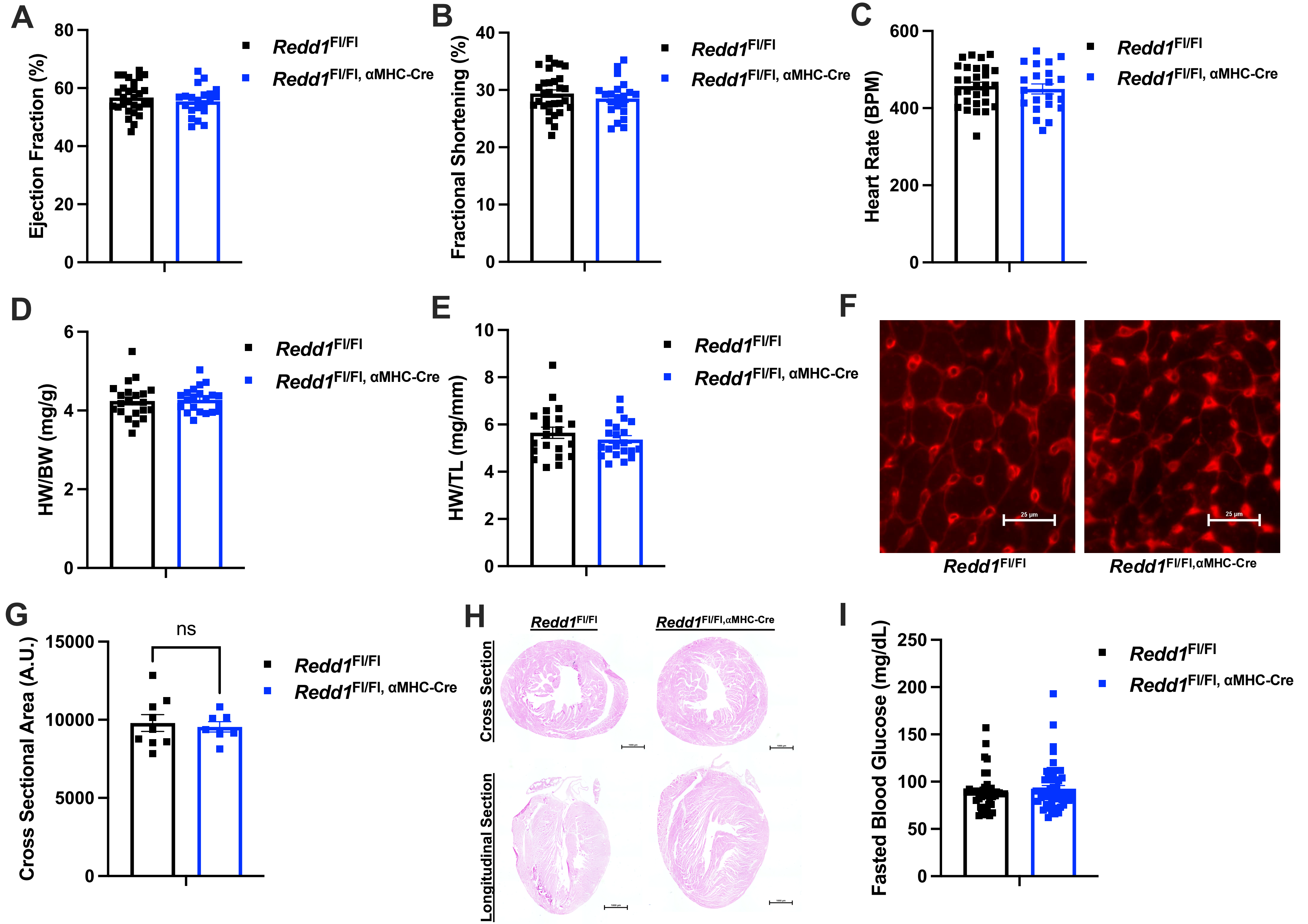

**Supplemental Figure 4. REDD1 deletion does not alter systolic function, cardiac size or structure, or fasted blood glucose.** Adult (10-14-week-old) male and female *Redd1*$^{Fl/Fl}$ and *Redd1*$^{Fl/Fl,\ \alpha MHC\text{-}Cre}$ mice were used. **A-C.** Mice were subjected to echocardiographic assessment with **(A)** ejection fraction, **(B)** fractional shortening, and **(C)** heart rate plotted. n=29,21 (ejection fraction, fractional shortening, and heart rate). **D-E.** Hearts were harvested and weighed. Heart weights (HW) were normalized to **(D)** body weight (BW) or **(E)** tibia length (TL), and plotted as a ratio (mg/g and mg/mm, respectively). **F-H.** Hearts were harvested, fixed, and stained with **(F)** 594 wheat germ agglutinin and **(G)** cardiomyocyte average cross-sectional area measured, n=9,7 or **(H)** hematoxylin and eosin. Representative images are shown. Scale bars are 25 µm and 1000 µm, respectively. **I.** After a 12 hour fast, blood glucose levels were assessed with a glucose meter. Concentrations are plotted as milligrams per deciliter. n=42,53.

Supplemental Figure 5.

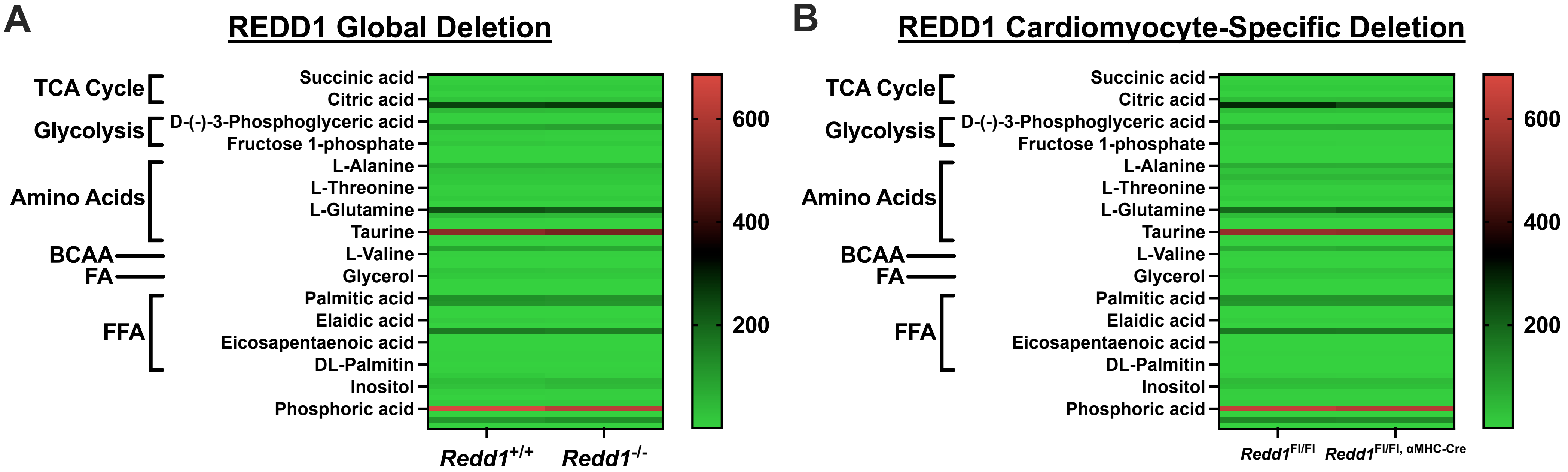

**Supplemental Figure 5. REDD1 deletion does not alter the cardiac metabolome.** Hearts of adult (8-14-week-old) male and female **(A)** *Redd1*$^{+/+}$ and *Redd1*$^{-/-}$ mice or **(B)** *Redd1*$^{Fl/Fl}$ and *Redd1*$^{Fl/Fl,\ \alpha MHC\text{-}Cre}$ mice were harvested and subjected to metabolomic profiling as described in the methods section.

### Supplemental Figure 6.

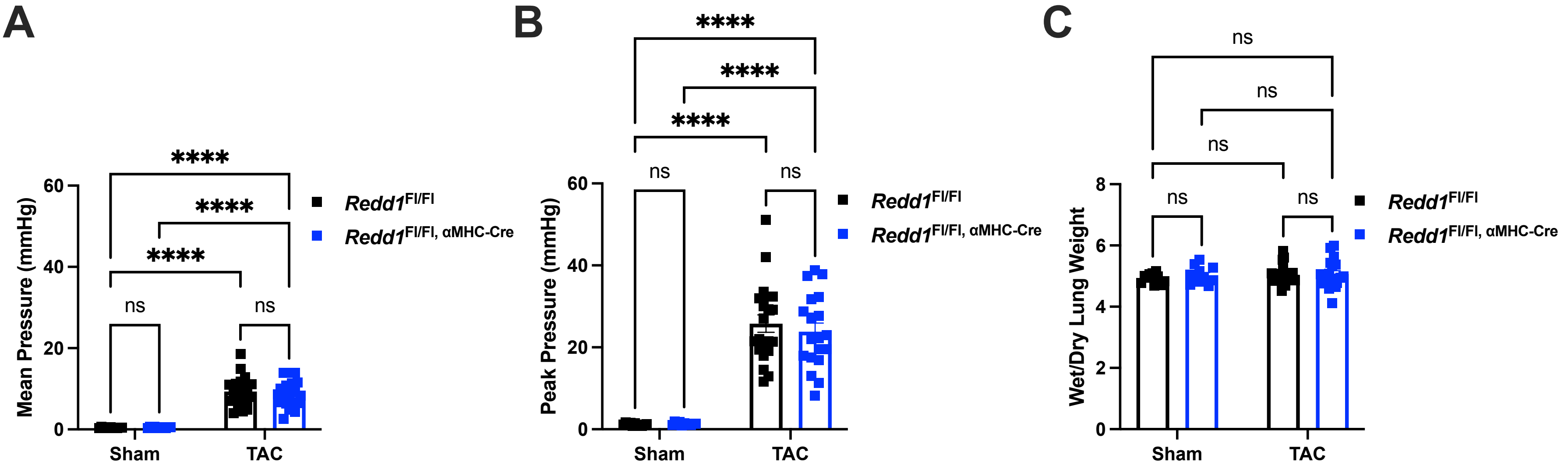

**Supplemental Figure 6. Two-week transverse aortic constriction (TAC) does not elicit heart failure.** Adult (10-12-week-old) male and female mice of the indicated genotypes were subjected to 2 weeks TAC or Sham operation. **A-B.** Doppler echocardiographic analyses were performed one-week post-operation and mean **(A)** and peak **(B)** aortic pressure gradients (mmHg) were calculated and plotted. n=11,11,21,19 (mean), n=11,11,21,19 (peak), 2-way ANOVA. **C.** Lungs were harvested and wet lung weights were normalized to dry lung weights and plotted. n=11,11,21,19, 2-way ANOVA. Error bars represent SEM. ****$p<0.0001$.

#### Supplemental Table 1.

| | *Redd1*$^{Fl/Fl}$ | *Redd1*$^{Fl/Fl,\alpha MHC\text{-}Cre}$ | p-value |
|---|---|---|---|
| **Heart Rate (BPM)** | 457.36 (±9.88) | 449.62 (±13.02) | 0.63 |
| **Diameter-S (mm)** | 2.76 (±0.07) | 2.86 (±0.06) | 0.27 |
| **Diameter-D (mm)** | 3.89 (±0.08) | 4.00 (±0.06) | 0.31 |
| **Volume-S (µL)** | 29.22 (±1.67) | 31.67 (±1.66) | 0.31 |
| **Volume-D (µL)** | 66.67 (±2.93) | 70.34 (±2.43) | 0.37 |
| **Stroke Volume (µL)** | 37.45 (±1.52) | 38.67 (±1.11) | 0.55 |
| **Ejection Fraction (%)** | 56.77 (±1.02) | 55.43 (±1.10) | 0.38 |
| **Fractional Shortening (%)** | 29.37 (±0.66) | 28.53 (±0.70) | 0.40 |
| **Cardiac Output (mL/min)** | 17.13 (±0.80) | 17.39 (±0.70) | 0.82 |

**Supplemental Table 1. Echocardiographic Assessment.** Hearts of adult (10-14-week-old) male and female *Redd1*$^{Fl/Fl}$ and *Redd1*$^{Fl/Fl,\ \alpha MHC\text{-}Cre}$ mice were subjected to echocardiographic assessment. Average values ± SEM for the indicated echocardiographic parameters are listed.
